# The SWI/SNF subunits BRG1 affects alternative splicing by changing RNA binding factor interactions with RNA

**DOI:** 10.1101/858852

**Authors:** Antoni Gañez Zapater, Sebastian D. Mackowiak, Yuan Guo, Antonio Jordan-Pla, Marc R. Friedländer, Neus Visa, Ann-Kristin Östlund Farrants

**Author notes:** Corresponding author: Ann-Kristin Östlund Farrants, Department of Molecular Biosciences, The Wenner-Gren Institute, The Arrhenius Laboratories F4, Stockholm University, SE 106 91 Stockholm, Sweden. Sebastian Mackowiak, Max Planck Institute for Molecular Genetics. Ihnestraße 63-73, 14195 Berlin, Germany. Antonio Jordan Plá, Departamento de Bioquímica y Biología Molecular, Facultad de Ciencies Biológicas, C/Dr.Moliner, 5,0, Burjassot 46100, Valencia University, Spain. Contributed equally (Sebastian D. Mackowiak, Yuan Guo).

## Abstract

BRG1 and BRM are ATPase core subunits of the human SWI/SNF chromatin remodelling complexes. The function of the SWI/SNF complexes in transcriptional initiation has been well studied, while a function in alternative splicing has only been studied for a few cases for BRM-containing SWI/SNF complexes. Here, we have expressed BRG1 in C33A cells, a BRG1 and BRM-deficient cell line, and we have analysed the effects on the transcriptome by RNA sequencing. We have shown that BRG1 expression affects the splicing of a subset of genes. For some, BRG1 expression favours exon inclusion and for others, exon skipping. Some of the changes in alternative splicing induced by BRG1 expression do not require the ATPase activity of BRG1. Among the exons regulated through an ATPase-independent mechanism, the included exons had signatures of high GC-content and lacked a positioned nucleosome at the exon. By investigating three genes in which the expression of either wild-type BRG1 or a BRG1-ATPase-deficient variant favoured exon inclusion, we showed that expression of the ATPases promotes the local recruitment of RNA binding factors to chromatin and RNA in a differential manner. The hnRNPL, hnRNPU and SAM68 proteins associated to chromatin in C33A cells expressing BRG1 or BRM, but their association with RNA varied. We propose that SWI/SNF can regulate alternative splicing by interacting with splicing-RNA binding factor and altering their binding to the nascent pre-mRNA, which changes RNP structure.

**Author summary:** Splicing, in particular alternative splicing, is a combinatorial process which involves splicing factor complexes and many RNA binding splicing regulatory proteins in different constellations. Most splicing events occur during transcription, which also makes the DNA sequence, the chromatin state and the transcription rate at the exons important components that influence the splicing outcome. We show here that the ATP-dependent chromatin remodelling complex SWI/SNF influences the interactions of splicing regulatory factors with RNA during transcription on certain exons that have a high GC-content. The splicing on this type of exon rely on the ATPase BRG1 and favour inclusion of alternative exons in an ATP-independent manner. SWI/SNF complexes are known to alter the chromatin structure at promoters in transcription initiation, and have been previously shown to alter the transcription rate or nucleosome position in splicing. Our results suggests a further mechanism for chromatin remodelling proteins in splicing: to change the interaction patterns of RNA binding splicing regulatory factors at alternative exons to alter the splicing outcome.

## Introduction

Chromatin influences transcription not only at the level of initiation and elongation: RNA processing is also influenced by the chromatin structure and changes are required to establish proper gene expression responses to the environment. Alternative splicing and alternative polyadenylation produce different mature mRNA from the same pre-mRNA, being an important source of the diversity of proteins (Wang et al., 2008; Di Giammartino et al., 2011). mRNA processing, such as 5’-capping, splicing and polyadenylation events occur to a large extent co-transcriptionally (Ameur et al., 2011; Tilgner et al., 2012), and is tightly coupled to the transcription machinery and chromatin (reviewed in Shukla and Oberdoerffer, 2012; Cusódio and Carmo-Fonseca 2015; Saldi et al., 2016). Processing factors and RNA binding factors are recruited by the RNA polymerase II (RNA pol II) and by chromatin during elongation. RNA pol II recruits factors by its C-terminal domain (CTD); the 5’-capping enzymes are recruited by serine 5-phophorylated (Ser5-P CTD) RNA pol II CTD (McCracken et al., 1997; Cho et al., 1997; Moteki and Price, 2002). Histone modifications in the gene body recruit chromatin proteins, such as the ATPase CHD1 which binds H3K4me3 at the start of a transcribed region and recruits U2snRNP (Sims et al., 2007). Histone-modifying and chromatin proteins are also recruited to the nascent RNA by endogenous small RNA bound to Argonaut (AGO) (Ameyar-Zazoua et al., 2012; Alló et al., 2014).

The mechanisms involved in the regulation of co-transcriptional alternative mRNA splicing are summarised in two general models: the *recruitment* and the *kinetic* model. The *recruitment* model proposes that the splicing outcome is a combinatorial event that depends on splicing factor recruited to the target exon. In addition to the general splicing machinery, many RNA-binding proteins, such as serine-rich proteins (SR-proteins) and heterogeneous nuclear ribonucleoproteins (hnRNPs) function as splicing enhancers and silencers (Witten and Ule, 2011; Deconte et al., 2013; Lee et al., 2015). These proteins bind to RNA and promote binding of the general splicing machinery or mask splice sites. The *kinetic* model postulates that the transcription rate determines the inclusion or skipping of alternative exons; a slow RNA polymerase II gives the splicing machinery more time to recognise splice sites and perform the splicing reaction (Kornblihtt, 2007; Kornblihtt et al., 2009; Ip et al., 2011; Naftelberg et al., 2015; Saldi et al., 2016). However, recent studies have shown that the transcription rate must be optimal to achieve a normal set of splice forms (Fong et al., 2017; Saldi et al., 2018). How the transcription rate is established and changed *in vivo* is poorly understood. It has been proposed that it depends on the phosphorylation state of the RNA pol II and on the modifications in the chromatin template. A higher Ser5-P CTD slows down or even pauses the RNA pol II, allowing for a time window for the splicing machinery to recognise weak splice sites (Batsché et al., 2006; Hirose et al., 2007; Harlen et al., 2016; Hsin and Manley, 2012; Costódio and Carmo-Fonseca, 2016; Garavis et al., 2017; Nojima et al., 2018). Furthermore, a number of histone modifications have been shown to localise with alternative exons and regulate transcription rate (Gundersen and Johnson, 2009; Luco et al., 2010; Hnilicova et al., 2011; Spain and Govind, 2011; Jonkers et al., 2014). The rate has also been associated with histone modifications that recruit different proteins, such as HP1α, which results in a slowdown of the RNA pol II and inclusion of exons (Chen et al., 2018; Iannone and Valcárcel, 2013; Zhou et al., 2014).

The *kinetic* model and the *recruitment* model are not mutually exclusive but rather potentiate each other, and many factors, such as CHD1, MRG15 and U2snRNPs, are recruited by histone modifications (Sims et al., 2007; Luco et al., 2010; Pradeepa et al., 2012; Yearim et al., 2015; Dujardin et al., 2014). Many of these adaptors that promote alternative splicing are chromatin proteins involved in chromatin dynamics. These proteins are usually part of chromatin remodelling complexes, and are important to establish specific chromatin states by altering the nucleosome occupancy (Hota and Bruneau, 2016; Clapier et al., 2017). It is well established that the ATP-dependent chromatin remodelling SWI/SNF complexes function at promoters, but the ATPases have also been implicated in different steps of RNA processing. The human SWI/SNF complexes are typically composed of either BRG1 or BRM as the ATPase catalytic subunit, and BAF155, BAF170 and Ini1/SNF5 (Hargreaves and Crabtree, 2013; Masliah-Planchon et al., 2015). The BRM SWI/SNF complexes have been proposed to change the phosphorylation state of the CTD of RNA pol II during elongation, which changes the rate of transcription (Batsché et al., 2006; Ito et al., 2008). The *Drosophila* SWI/SNF complex changes the splicing outcome of a number of transcripts (Tyagi et al., 2009; Waldholm et al., 2011), by nucleosome stability (Zraly and Dingwall, 2012). SWI/SNF ATPases have also been found to associate to the growing RNP (Tyagi et al., 2009) and to interact with general splicing factors (Zhao et al., 1998; Dellaire et al., 2002; Ito et al., 2008; Allemand et al., 2016; Yu et al., 2018). Furthermore, the human BRG1 regulates alternative cleavage sites by degrading the 3’ end processing factor CstF through interacting with BRCA/BARD (Fontana et al., 2017). BRG1 and its *Drosophila* orthologue Brm are also involved in cleavage site choice of mRNA by interacting with members of the cleavage and polyadenylation factor complexes (CPSF) (Yu et al., 2018). However, the mechanisms by which the SWI/SNF ATPases function in alternative mRNA processing is still not fully understood.

In this study we have performed an RNA-seq transcriptome analysis of C33A cells, a SWI/SNF deficient cell line (Muchardt and Yaniv 1993; Wong et al., 2000; Decristofaro et al., 2001), that exogenously expresses SWI/SNF ATPases, and we have identified a subset of genes whose splicing outcome was affected. Both exon inclusion and skipping of exons were favoured by expression of the ATPases, and approximately half did not require the ATPase activity. BRG1 affected exons separated into two groups, ATPase dependent and ATPase independent, of which the ATPase independent exons exhibited specific features: a high GC-content without a nucleosome positioning at the exon. We show that for this group of exons, the effect of BRG1 on splicing is not correlated with an altered nucleosome density or change in RNA pol II accumulation. We show instead that BRG1 can rearrange the binding of RNA regulators to RNA, and thereby fine-tune the splicing outcome.

## Results

### BRG1 and BRM affect the splicing outcome of a subset of genes

SWI/SNF ATPases and complexes have been shown to affect splicing in both human cell lines and in *Drosophila* (Batshé et al., 2006; Ito et al., 2008; Tyagi et al., 2009; Allemand et al., 2016), but the effect has only been shown on a few genes. To identify the extent of the effect of SWI/SNF ATPases in splicing, we performed RNA-seq of the polyadenylated transcriptome of C33A cells transfected with either BRG1 or BRM. C33A cells transfected with empty vector was used as reference, and duplicates of BRG1 or BRM transfected cells, as well as cells transfected with the ATP deficient variants, were performed. After 48 hours of transfection, RNA was prepared and converted to cDNA using oligo-dT primers. The results from the RNA seq data were analysed for differentially spliced exons in the ATPase expressing cells, using the MISO algorithm (Katz et al., 2010), and a subset of genes with differentially spliced exons was identified (Supplementary Table S1). Both BRG1 and BRM affected the splicing outcome; BRG1 expression resulted in altered splicing of 836 exons (Figure 1A), of which 56% exhibited an increased inclusion (Figure S1A). Expression of the ATPase deficient BRG1 (BRG1-mut) also affected splicing: 1052 exons were affected (Figure 1A), with 57% exhibiting an increased inclusion (Figure S1A). The ATPase activity was not required for the splicing outcome in 38% of the BRG1 affected exons; 316 exons were common to both BRG1 and BRG1-mut, whereas 520 exons were influence by BRG1 alone (Figure 1A). Of the 836 exons affected by BRG1, 240 were also affected by the expression of BRM (Figure 1A), suggesting that most BRG1 target exons are exclusive to BRG1. BRM expression affected more exons: 1116 exons, 49% of which exhibited favoured inclusion. Only a small number, 195, of these were ATPase independent, and appeared in cells expressing BRM-mut (Figures S1A and S1B).

**Figure 1.**
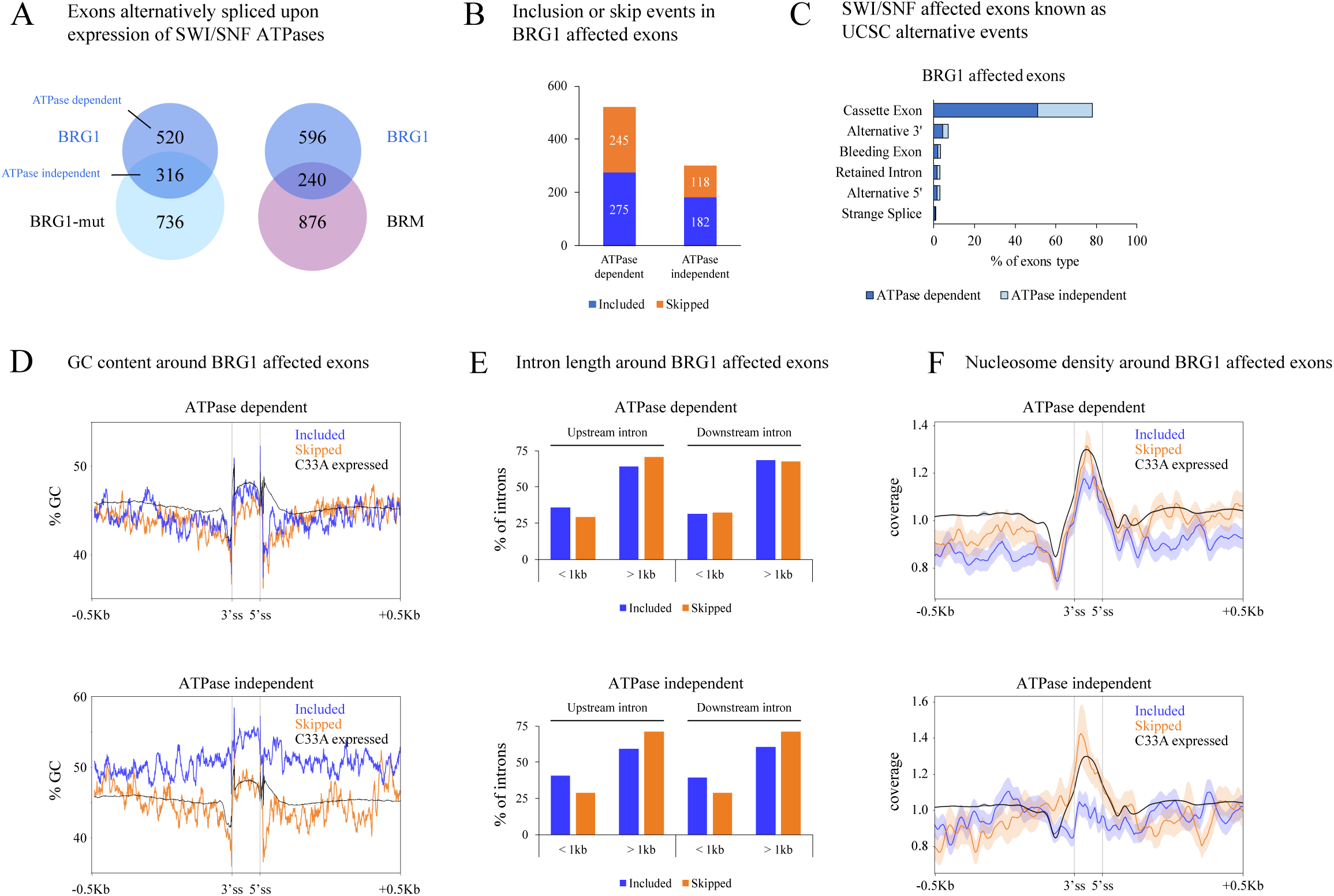
BRG1 and BRM affect alternative splicing of a subset of genes. **A)** Venn diagram showing exons affected by the exogenous expression of BRG1-wt, BRG1-mut and BRM-wt. Exons affected only by BRG1-wt are referred to as “BRG1 ATPase dependent” while exons affected by both BRG1-wt and BRG1-mut are referred to as “BRG1 ATPase independent”. pOPRSVI plasmid was used as control. RNA was harvested 48 h after transfection and measured by RNA-seq, where differential exon levels were estimated with the MISO algorithm (Katz et al., 2010). **B)** Number of exons with increased inclusion (blue) or skipped (orange) upon expression of BRG1. **C)** UCSC alternative events classification of BRG1 affected exons detected by MISO, represented as percentage of the total amount of affected exons detected. **D)** GC content in ATPase dependent (top panel) and ATPase independent (bottom panel) BRG1 affected exons +/- 500bp harbouring regions divided into included (blue) and skipped (orange) exons. Exons are plotted as 100 bp, each bp representing the average GC content of the 1% of the total length for each exon. Exons and the harbouring +/-500 bp regions show the 10% of GC in each position. **E)** Length of upstream and downstream introns from ATPase dependent (top panel) and ATPase independent (bottom panel) BRG1 affected exons. Bars represent the percentage of affected exons shorter or longer than 1 kb. **F)** Nucleosome density from ATPase dependent (top panel) and ATPase independent (bottom panel) BRG1 affected exons, generated with DeepTools2 (Ramírez et al., 2016) from publicly available data from the K562 cell line, at the exons differentially included (blue) or skipped (orange) by the SWI/SNF ATPases and the +/- 0.5 Kb harbouring region (light colour shows standard error).

Next, we analysed whether the genes with the affected exons were differentially expressed, since the primary action of SWI/SNF complexes is to regulate chromatin at promoters and enhancers. Analysis performed using DESeq2 (Supplementary Table S2) showed that the expressions of 251 genes were affected upon expression of BRG1 in C33A cells, and the genes affected showed a great overlap with the genes found in BRM expressing cells (Figure S1C). Interestingly, BRG1-mut expression also affected gene expression, which indicates that genes also in human cells genes are regulated by SWI/SNF in an ATPase independent manner, similar to *Drosophila* cells (Jordán-Pla et al., 2018) (Supplementary Table S3). Only a minor fraction, 84 genes, were affected at the levels of both gene expression and splicing, suggesting that SWI/SNF complexes in most cases affect the promoter and exons independently.

Most studies have focused on the function of BRM in RNA processing, and the role of BRG1 in splicing is poorly understood. Our results show that BRG1 also affects splicing, which prompted us to focus on role of the BRG1-SWI/SNF in splicing. The expression of BRG1 or BRG1-mut did not affect exons that overlapped completely, and we divided them into two groups for further analysis. We argued that the group of exons affected by expression of either BRG1 or BRG1-mut used an ATP-independent mechanism, whereas those exons affected by BRG1 alone used an ATPase dependent mechanism. Both groups contained included and skipped exons; 53% included exons in the BRG1 ATPase dependent group and 55% included exons in group of ATPase independent affected exons (Figure 1B). To examine whether SWI/SNF subunits associated with the affected genes, we analysed published ChIP-seq data from HeLa cells of BRG1 and several SWI/SNF core subunits (Euskirchen et al., 2011). The factors BRG1, SNF5/INI1, BAF155 and BAF170 associate with the genes harbouring differentially spliced exons in the affected ATPase dependent and independent groups (Supplementary Figure S1D).

The overlap in signals suggested that BRG1 associated with the affected exons in the context of a SWI/SNF complex and not as a single ATPase. We also analysed the binding to the genes containing BRG1 affected exons of other DNA binding proteins in the OREGANNO database (Lesurf et al., 2016). BRG1 (SMARCA4 gene product) was the most abundant protein present at the genes with exons affected with BRG1, both in the ATPase dependent and independent way, and other factors, such as EGR1 and CTCF, also bound to these genes (Supplementary Figure S1E).

### BRG1 included and skipped exons have different GC content and chromatin signatures

The MISO analysis identifies internal exons and we established that the majority of the BRG1 affected exons were cassette exons (80%) (Figure 1C). Exons have been classified depending on different features, such as sequence characteristics, chromatin states (Hollande et al., 2018) and the association of splicing enhancer and silencing proteins (Lee and Rio, 2015). We thus examined the differentially spliced exons in the BRG1 ATPase dependent and ATPase independent groups for specific features. The GC content of exons is one signature that has been proposed to be involved in exon and intron definition mechanisms (Zhang et al., 2011; Amit et al., 2012). The affected included and skipped exons in the BRG1 ATPase independent group exhibited different GC contents, and the ATPase independent exons included a high GC content (approximately 55%), flanked by relatively high GC content introns (+/– 500 bp) (Figure 1D). In contrast, the skipped exons had a lower GC content (47%) similar to the mean of all expressed exons (Figure 1D). The GC content was low, at the level of the mean of all expressed exons, in both included and skipped exons in the ATPase dependent exons (Figure 1D).

Exons with a high GC content are usually flanked by short introns, which favour intron definition mechanisms in which splice sites are defined by the binding of general splicing factors (Zhang et al., 2011; Amit et al., 2012; Georgomanolis et al., 2016). Most of the introns surrounding included exons both upstream and downstream in the two groups were longer than 1 kb: 59% of the exons affected in ATPase dependent group and 64% of exons in the ATP independent group (Figure 1E). This suggests that the length of the introns is not a defining characteristic for BRG1 affected exons.

Since positioned nucleosomes are also exon or intron definition features (Kornblitt et al., 2009; Schwartz and Ast, 2010; Amit et al., 2012), the BRG1 affected exons were examined for nucleosome occupancy using ENCODE data. The BRG1 ATPase dependent affected exons had positioned nucleosomes in both skipped and included exons (Figure 1E). The nucleosome occupancy in the BRG1 ATPase independent group displayed a pattern in which skipped exons had a nucleosome positioned at the exon, while no such positioned nucleosome was observed at included high GC-content exons (Figure 1F). We also investigated whether BRG1 affected exons that were affected by BRM displayed differences between included and skipped exons. 122 of the 240 BRG1 and BRM common exons were also affected in BRG1-mut expressing cells (Supplementary Figure S1B) and when these were removed, both included and skipped BRG1 and BRM exons had a low GC-content and a positioned nucleosome at the exon (Supplementary Figure S1F). Since GC-content and nucleosome positioning are features that have been proposed to determine splicing mechanism, we suggest that these underlying features also influence the mechanisms used by BRG1; ATPase dependent included and skipped exons use a positioned nucleosome to define the exon in a low GC environment, whereas included exons with a high GC-content are ATPase independent and defined by a different mechanism.

Several reports have suggested that exons, also alternative exons, hold specific combinations of histone modifications, such as H3K4me3, H3K9Ac, H3K27Ac, and H3K36me3 (Enroth et al., 2012; Iannone and Valcárcel, 2013; Curado et al., 2015; Hou et al., 2017; Kim et al., 2018), which led us to analyse the chromatin state at the BRG1 affected exons using the ENCODE data. BRG1 affected exons had slightly more exons exhibiting a chromatin state of transcriptional elongation than the average in all expressed exons (Figure S1G). The ATPase independent exons also had slightly more exons in a transcription start site configuration (Figure S1G). We also analysed differences in histone modifications in the ATP dependent and independent affected exons and their flanking region (+/- 2 kbp from the 3’ and 5’ splice sites) using ENCODE data. The ATP independent included exons had a higher level of H3K27Ac than ATP dependent, while the ATPase dependent exons had a lower level of H3K36me3 ATPase dependent skipped exons (Supplementary Figure S1H). The H3K4me3 and H3K9ac patterns of the ATPase dependent and the independent exons displayed were similar (Supplementary Figure S1H).

### MYL6, GADD45A and MAZ are genes that are alternatively spliced by BRG1 and BRM

The ATPase independent BRG1 included exons exhibit high GC-content without a nucleosome positioned at the exon, a group of exons described by Amit et al. (2012). This group of exons was proposed to use exon definition mechanisms instead of the more commonly used intron definition mechanisms (Amit et al., 2012). We chose to further elucidate the mechanism by which BRG1-SWI/SNF complexes affect splicing on the inclusion of high GC-content alternative exons. Three genes with increased inclusion in C33A cells expressing BRG1 were selected for further investigation: MYL6, GADD45A and MAZ. These had been identified through our genome-wide MISO analysis (Supplementary Table S1) as having a higher level of a cassette exon included in the ATPase independent group.

Two alternatively spliced forms of MYL6 were expressed in C33A cells; the one with the cassette exon 6 included was less abundant, 13% of the expressed transcripts (Figure 2A, bottom panel). The abundances of the isoforms, determined by PCR, were similar to those observed in the RNA-seq, which showed that 8.5% constituted the MYL6 isoform including exon 6. qPCR analysis (using primers in Figure 2A, top panel) validated a higher inclusion level of exon 6 by BRG1, BRG1-mut and BRM expression (Figure 2A, middle panel) with over 20% above the level observed in control cells. GADD45A exon 2 was identified as being more included by BRG1 in the MISO analysis, but the splice form was the most abundant one in C33A, with 78% of isoform containing exon 2 in C33A cells (75% in the RNA seq analysis) (Figure 2B, bottom panel). qPCR analysis showed that expression of the BRG1, BRG1-mut and BRM increased the inclusion of exon 2 by between 40% and 60% (Figure 2C, middle panel). The MISO analysis also showed a favoured inclusion of exon 5 in the MAZ transcript, which was the low abundant form, only constituting 19%, in C33A cells (8% in the RNA seq analysis) (Figure 2C, bottom panel). qPCR analysis showed that exon 5 was significantly more included upon BRG1 and BRM expression when compared to the transcript without exon 5, by 20% and 40%, respectively (Figure 2C, middle panel). We conclude that these exons were affected by both BRG1 and BRM, and neither of them changed the ratio between affected exons dramatically, but rather fine-tuned the abundance of the different transcripts.

**Figure 2.**
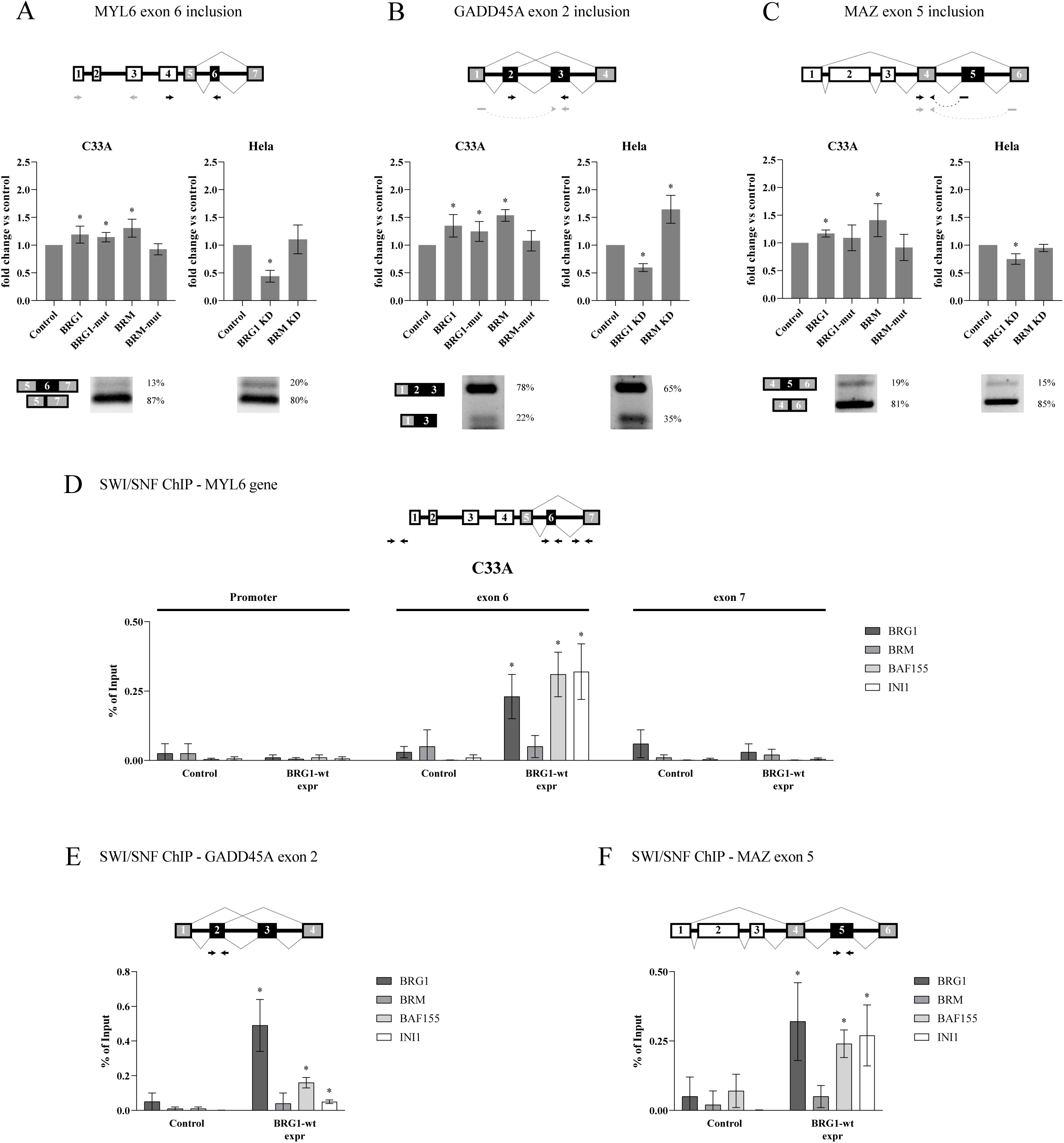
Validation of MYL6, GADD45A and MAZ genes. **A-C)** Top panels show the scheme of the affected exon in the gene context showing the affected exon (black), exons taken as reference by MISO to calculate differential inclusion (grey), and constitutive exons (white); arrows show the position of primers used in qPCR with one pair targeting the affected exon (black) and another pair used for normalisation (grey). Middle panels show the differential exon inclusion upon exogenous expression of SWI/SNF ATPases in C33A (left) or knock-down in Hela (right) measured by qPCR; asterisks denote significant differences compared to the control (p-value < 0.05), n = 5. Bottom panels show the relative basal levels of the two isoforms with included or skipped exons detected by MISO, in control cells in C33A (left) and Hela cells (right). **D-F)** ChIP-qPCR shows the association of BRG1, BRM, BAF155 and INI1 at the MYL6 gene in the promoter region, affected exon 6 and constitutive exon 7 (D), GADD45A exon 2 (E) and MAZ exon 5 (F). The association is presented as percentage of input (n = 6), and significant changes (p-value < 0.05) compared to control are marked with asterisks. Top panels show the positions of the primers used for ChIP-qPCR targeting the promoter region, affected exon 6 and constitutive exon 7 from the MYL6 gene (A), and affected exons from GADD45A (B) and MAZ (C).

We also measured the level of splicing of these exons upon knock-down of BRG1 or BRM in HeLa cells, which express both ATPases endogenously. The endogenous pattern of expression of MYL6 exon 6 splice variants showed that the longer variant was the less abundant, with approximately 20% of the transcript. BRG1 knock-down in HeLa cells led to a reduced inclusion of the exon by 50%, while BRM knock-down did not lead to a difference in the inclusion rate (Figures 2A, middle panel). The longer splice variant of GADD45A in HeLa cells was the most prominent form, 65% (Figure 2B, bottom panel). BRG1 knock-down in HeLa cells resulted in the longer splice variant being less abundant (Figure 2D, middle panel), whereas knock-down of BRM resulted in a further increase in inclusion of exon 2 (Figure 2B, middle panel). The longer splice variant of MAZ constituted 15% of the mRNA in HeLa cells (Figure 2C, bottom panel). Knock-down of BRG1 in HeLa cells reduced the inclusion of exon 5, and similar to exon 6 in MYL6, no effect was observed in BRM knock-down cells (Figure 2C, middle panel). We conclude that the two ATPases are not able to fully substitute for each other and this effect is cell type specific.

### BRG1 affects splicing as part of the SWI/SNF complex

The results from the qPCR analysis showed that the actions of BRG1 and BRM are different on the three genes in both C33A and HeLa cells. In C33A cells, expression of BRG1 and BRG1- mut affected splicing, while only BRM and not BRM-mut expression altered the splicing pattern. We conclude that BRG1 favoured inclusion of these cassette exons in an ATP-independent manner, whereas BRM favoured inclusion in ATP dependent manner. To further investigate the mechanisms by which SWI/SNF complexes influence splicing, we focused on the BRG1 ATPase independent action.

We analysed by ChIP-qPCR whether the exogenously expressed BRG1 in C33A was recruited to the affected exon in the three genes investigated. BRG1 was recruited to the exons of MYL6, GADD45A and MAZ in BRG1 expressing cells, while the BRM levels were low (Figures 2D, 2E and 2F). BAF155 and INI1/hSNF5, core subunits of SWI/SNF complexes, were also present at the exons in BRG1 expressing cells but not in control cells (Figure 2D), which supports the idea that the BRG1 functions as part of the SWI/SNF complex. The BRG1 or the core subunits were not recruited to the promoter or the constitutive exon 7 of MYL6 (Figure 2D), showing that the association of SWI/SNF subunits was specific to the affected exon. BRG1-mut was also detected at the affected exons in cells expressing BRG1-mut (Supplementary Figure S2A). In addition, we examined the association of BRM to the exons in the BRM and BRM-mut expressing cells, and found that these ATPases were also recruited to these exons (Supplementary Figure S2B). In the case of BRM-mut, this did not affect the splicing outcome.

### Expression of BRG1 does not change the chromatin landscape

SWI/SNF complex components in *Drosophila* cells have been shown to change nucleosome configuration to achieve splicing differences (Zraly and Dingwall. 2012). Furthermore, in mammalian cells, SWI/SNF complexes are suggested to alter the RNA polymerase rate and phosphorylation level to favour inclusion of exons (Batshé et al., 2006; Ito et al., 2008). This led us to investigate whether BRG1 and BRG1-mut expression in C33A cells induces changes in the nucleosome density over the exons. We performed ChIP of histone H3 and of several histone modifications, but none of the changes observed correlated with splicing outcome. The histone H3 level at MYL6 exon 6 remained at the same level in control cells and in cells that expressed BRG1, while it was reduced by the expression of BRG1-mut (Figure 3A). This pattern was also observed at the constitutive exon 7, which was not BRG1 or BRG1-mut dependent. The histone H3 occupancy at GADD45A resembled that at MYL6 exon 6 (Figure 3B), and BRG1-mut reduced the level. No change in occupancy was observed on MAZ exon 5 (Figure 3C). The occupancy in BRM and BRM-mut expressing cells was altered compared to control cells on MYL6 and MAZ exon 5, but not on GADD45 exon 2 (Supplementary Figures S3A to S3C). We conclude that the nucleosome landscape does not affect splicing on these genes.

**Figure 3.**
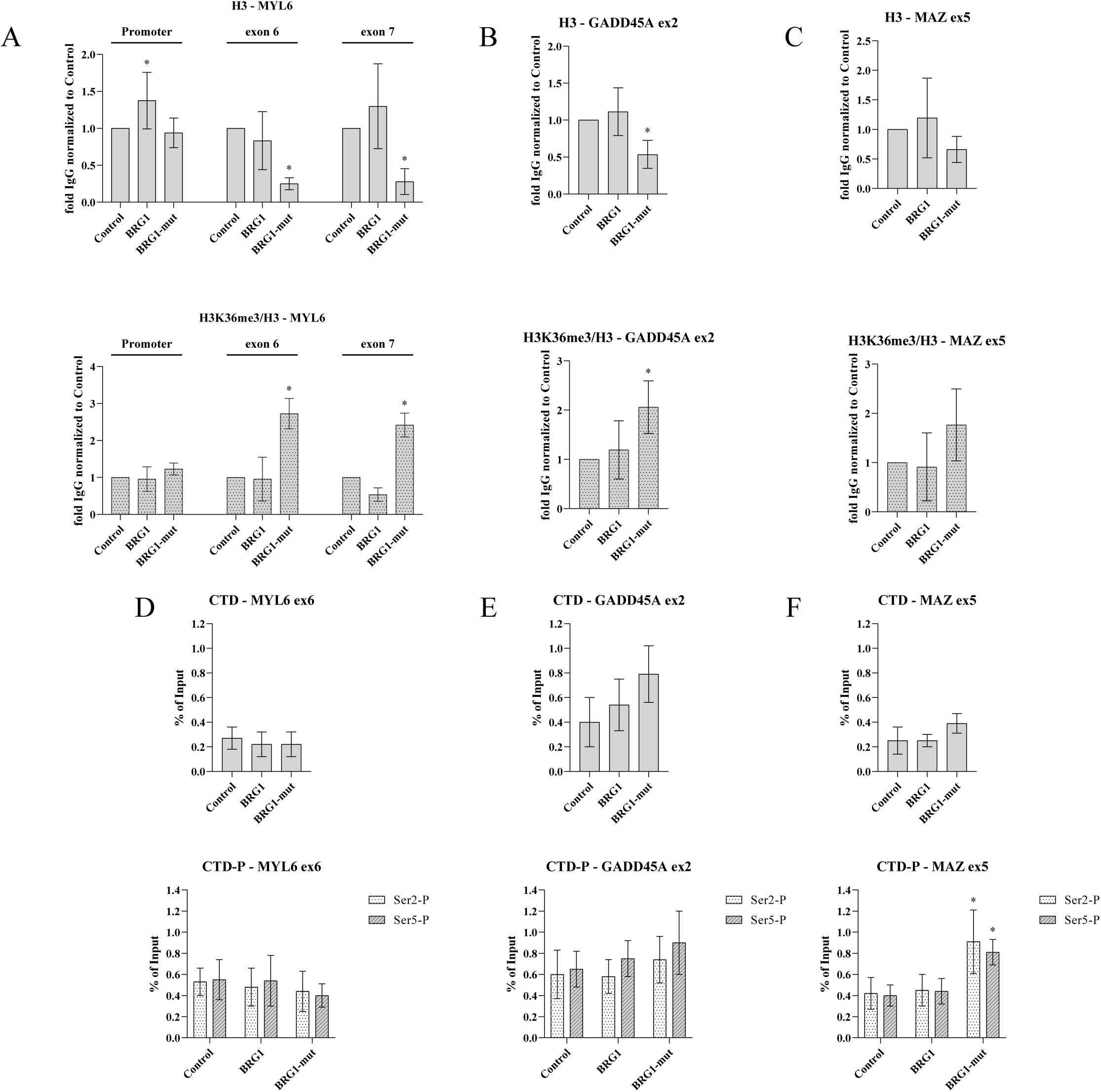
Nucleosome and polymerase density in the affected exons. **A-C)** ChIP-qPCR using antibodies against H3 (top panels) and H3K36me3 (bottom panels) targeting MYL6 gene (A), GADD45A exon 2 (B) and MAZ exon 5 (C) in C33A cells expressing BRG1 and BRG1-mut. H3K36me3 levels were normalized to H3. The association is related to the association in control cells, and significantly different values (p-value < 0.05) are denoted by asterisks (n = 3). **D-F)** ChIP-qPCR using antibodies against polymerase II CTD (top panels), and phosphorylated serines 2 and 5 from polymenrase II CTD (P-CTD, bottom panels) targeting MYL6 gene (A), GADD45A exon 2 (B) and MAZ exon 5 (C) in C33A cells expressing BRG1 and BRG1-mut. The association is related to the association in control cells, and significantly different values (p-value < 0.05) are denoted by asterisks (n = 5).

We also investigated the accumulation of the histone modifications H3K36me3, H3K4me3, H3K9Ac and H3K27Ac, but no changes that could be correlated with the splicing outcome were observed (Figures 3A to 3C, bottom panel, and Supplementary Figures S3A to S3F). BRG1-mut expressing cells displayed a slight increase in H3K36me3 in MYL6 exon 6 and exon 7 as well as in GADD45A exon 2 (Figures 3D to 3B). Histone H3K4me3 changed in BRG1-expressing cells, higher at MYL6 exon 6 and GADD45A exon 2, and lower levels at MAZ exon 5. The levels of these modifications in BRM and BRM-mut expressing cells varied, and both accumulation and reduction in H3K36me3 and H3K4me3 were observed at the different genes (Supplementary Figures S3A toS3F). However, no changes in histone density or histone modifications correlated with changes in splicing outcome, and we conclude that BRG1 does not affect the chromatin landscape around affected exons to promote inclusion of this group of exons.

### Expression of the SWI/SNF ATPases does not change the RNA pol II occupancy

Next, we examined the effect of the expression of BRG1 and BRG1-mut on the RNA pol II occupancy at the investigated exons. The occupancy was not significantly changed at any of the exons in the three genes compared to that in control cells (Figures 3D to 3F and Supplementary Figures S3D to S3I). Only BRM-mut expressing cells displayed a reduced RNA pol II CTD accumulation in exon 5 in MAZ compared to control cells (Supplementary Figure 3I). The phosphorylation of CTD at serine 2 (Ser2-P CTD), and in particular at serine 5 (Ser5-P CTD), plays an important role in the dynamics of RNA pol II and splicing (Batsché et al., 2006; Ito et al., 2008; Ip et al., 2011; Nojima et al., 2018). No differences were found in the levels of Ser2-P CTD or Ser5-P CTD at the exons of MYL6 and GADD45A when expressing BRG1 and BRG1-mut compared to control cells (Figures 3D and 3E, lower panel). However, MAZ exon 5 displayed an increase of both Ser2-P CTD and Ser5-P CTD in BRG1-mut-expressing cells (Figure 3H), indicating that BRG1-mut recruitment in the context of MAZ exon results in altered RNA pol II phosphorylation levels. No changes in RNA Pol II phosphorylation were observed in BRM and BRM-mut expressing cells (Supplementary Figures S3G to S3I). In summary, the expected increase in the Ser5-P CTD to Ser2-P CTD ratio, or even a consistent accumulation of phosphorylated RNA pol II CTD, failed to occur on the investigated included exons in BRG1 expressing cells. We therefore conclude that BRG1-SWI/SNF complexes are able to influence the splicing outcome in alternative ways than by changing chromatin landscape or RNA pol II CTD state. We hypothesise that BRG1-SWI/SNF complexes instead may alter the recruitment of regulatory RNA binding proteins to influence the splicing outcome in an ATP-independent way on this group of exons.

### BRG1 and BRM interact with RNA binding proteins, including splicing regulatory proteins

Exons with high GC-content flanked by introns with high GC-content have been proposed to favour intron definition mechanisms in which splice sites are defined by the binding of splicing factors (Amit et al., 2012; Georgomanolis et al., 2016). SWI/SNF ATPases, in particular BRG1, interact with several RNA binding proteins and general splicing factors (Zhao et al., 1998; Dellaire et al., 2002; Tyagi et al., 2009; Allemand et al., 2016). We analysed the mass spectrometry data of BRG1 and BRM interacting proteins in the RNAse treated chromatin fraction (RNP fraction) from HeLa cells (Yu et al., 2018). Several RNA binding proteins were found in the co-immunoprecipitate; an enrichment analysis for the GO term “RNA binding” (GO:0008380) revealed 79 interactors with BRG1 and 18 interactors with BRM, 16 of them identical in the two groups (Figure 4A). The top significant GO terms from BRG1 and BRM interactors include “mRNA splicing, via spliceosome” and “spliceosome complex” (Supplementary Figure S4A), strongly suggesting a close relation between SWI/SNF ATPases and the splicing machinery. We compared the interactors associated with BRG1 and BRM in the RNP-fraction to defined splicing factors and RNA binding factors (Hegele et al., 2012) and the results showed that BRG1 and BRM interacted with factors involved in different steps in the splicing cycle (Figure 4B). BRG1, and to some extent BRM, interacted mainly with the peripheral protein complexes that are recruited during the assembly of the A to Bact/B* complexes, such as the 3’-splice recognition proteins U2AF2 and SF1 in the A complex and the general splicing factors in the U2 snRNP complexes SF3a and SF3b, the U4/U6 snRNP factors Prp31 and Prp3, U5 snRNA component BRR2, Prp8 and Prp6, and the Prp19/CDC5Lcomplex in the Bact/B*. In addition to factors recruited early, proteins involved in later steps, such as the release of the spliceosome by Prp43/DHX15, as well as proteins of the exon junction complex and the THOC complex (EJC/TREX) were also found to bind to BRG1. Splicing regulators were also found; hnRNP proteins constituted a large group that interacted with BRG1 and BRM. We validated the interactions of proteins representing different groups in the mass spectrometry analysis: the RNA binding factors hnRNPL, DHX9, THOC2, the U2 splice factor SAP155 and the alternative splicing regulator Sam68 (Supplementary Figures S4B and S4C).

**Figure 4.**
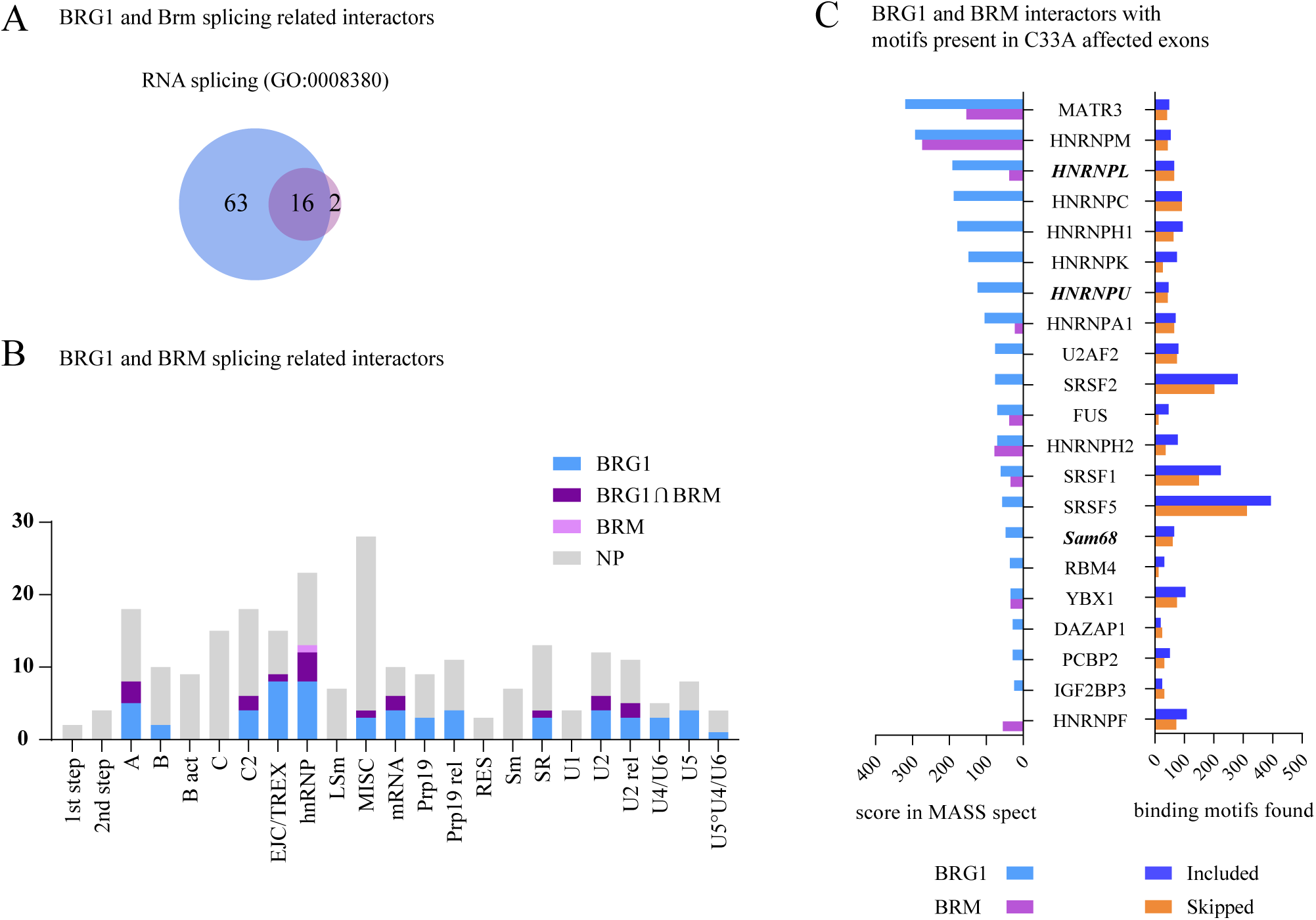
SWI/SNF interactions with RNA binding factor. **A)** Venn diagram with the number of RNPs interacting with BRG1 and BRM in the nascent RNA (data from Yu et al., 2018), that are associated with the GO term RNA splicing (GO:0008380). **B)** Number of BRG1 and BRM interacting proteins extracted from Yu et al. (2018), known to be splicing (classification from Hegele et al., 2012). The total length of the bar corresponds to the total amount of proteins classified in each group. BRG1 interacting protein in blue, BRM in pink, both present in BRG1 and BRM in purple, and not present (NP) in any IP in grey. **C)** RNA binding proteins interacting with BRG1 or BRM ordered by their score from MASS-spectroscopy (left) with known motifs present in affected exons (right) classified into included (blue) or skipped (orange) exons. The RNA binding proteins selected for detailed mechanistic studies (hnRNPL, hnRNPU and Sam68) are depicted in bold-italics.

Binding motifs of several RNA binding factors were present in the exons affected by BRG1 expression or in the immediate flanking regions (Figure 4C, right lane). We compared factors whose motifs were found at the exons with the factors found as interactors with BRG1 and BRM in the mass spectrometry (Figure 4C, left lane) and found that many hnRNPs were well represented. They had both binding motifs and were found to bind to the ATPases in the mass spectroscopy analyses with high scores.

### BRG1 recruits RNA binding factors to the affected genes

Next, we asked whether the effect on the splicing outcome observed upon expression of BRG1 was caused by recruitment of RNA binding proteins to the exons investigated. By ChIP, we showed that expression of BRG1 and BRG1-mut changed the pattern of factors associating with MYL6 exon 6, and a number of factors were recruited, such as hnRNPL, hnRNPU, hnRNPA1, hnRNPA2B1, DHX15, SYF1 and SAM68, although to different levels (Figure 5A). HnRNPU was already present in control cells and remained associated with the site in BRG1 expressing cells. Expression of BRM, and particularly BRM-mut, recruited fewer factors to the exons than BRG1 (Supplementary Figure S5A). We focused on hnRNPL, hnRNPU and SAM68 for further investigations, since hnRNPL strongly associated with BRG1 in the immunoprecipitation of the RNP fraction, hnRNPU is reported to bind BRG1 and histone modifying proteins (Obrdlik et al., 2008; Vizlin-Hodzic et al., 2011), and SAM68 regulates splicing together with BRM (Batsché et al., 2006). Of the three factors, SAM68 was not recruited to MYL6 exon 6 by the expression of BRG1 (Figures 5A and 5B) or BRG1-mut (Figure 5A), but by BRM (Supplementary Figure S1B). To examine the specificity of the recruitment to the exon, we also examined the promoter and the constitutive exon 7, which harbours a potential hnRNPL site (Hung et al., 2008; Rossbach et al., 2012), for association of these factors. No factor associated with the promoter upon expression of BRG1 and SAM68 associated with the constitutive exon 7, either in control cells or in BRG1 expressing cells (Figure 5B), suggesting that BRG1 altered the factor binding at the affected exons. The recruitment of factors by BRM was also specific to exon 6, and the promoter and exon 7 exhibited the same pattern of factors as that found in BRG1 expressing cells (Supplementary Figure S5B). This suggests that the ATPases recruit factors directly to the alternative exon 6 without affecting the exons in the vicinity. However, the pattern of factors recruited depended on the ATPase expressed. Interestingly, even BRM-mut recruited several factors (Supplementary Figure S5A), although it did not alter the splicing outcome. This prompted us to investigate whether BRG1 not only recruited splicing regulators to chromatin, but also affected their binding to RNA.

**Figure 5.**
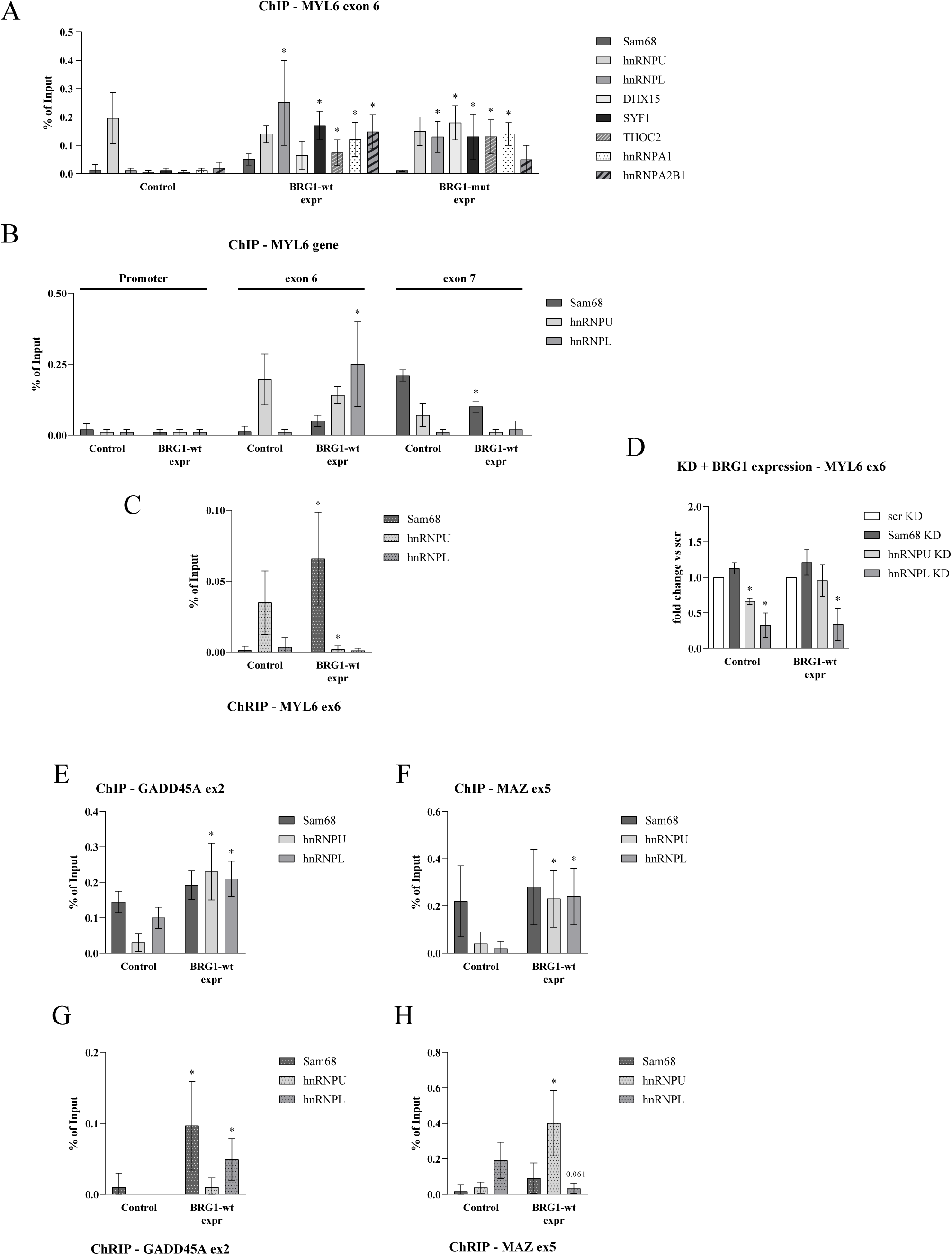
RNA binding factor are recruited by BRG1 and BRM to the target sites. **A)** ChIP-qPCR targeting MYL6 exon 6 in C33A cells expressing BRG1-wt and BRG1-mut was performed with antibodies against Sam68, hnRNPU, hnRNPL, DHX15, SYF1, THOC2, hnRNPA1 and hnRNP2B1; significant changes (p-value < 0.05) compared to control are marked with asterisks. **B)** ChIP was performed with antibodies against Sam68, hnRNPU, hnRNPL and analysed with qPCR with the same primers as in Figure 2 for MYL6 promoter, exon 6 and exon 7 in cells expressing BRG1-wt. Results are presented as percentage of input, and asterisks show significant changes (p-value < 0.05) compared to control (n = 6). **C)** ChRIP was performed with antibodies against Sam68, hnRNPU, and hnRNPL, and analysed with qPCR with the same primers as in Figure 2 for MYL6 exon 6 in cells expressing BRG1-wt. Results are presented as percentages, and asterisks show significant changes (p-value < 0.05) compared to control (n = 4). **D)** Changes in the inclusion of SWI/SNF affected exons after knocking down Sam68, hnRNPU and hnNRPL. C33A cells were transfected with siRNA that targeted RNA binding factors interacting with BRG1 and BRM, incubated for 24 h and transfected with BRG1-wt expressing for additional 48 h. The primers presented in Figure 2A were used to assess exon 6 in MYL6. Values were normalised to scr siRNA for each ATPase expressed and asterisks show significant changes (p-value < 0.05) compared to control (n = 3). **E-F)** ChIP was performed in C33A cells expressing BRG1-wt with antibodies against Sam68, hnRNPU, and hnRNPL, and analysed with qPCR with the same primers as in Figure 2 for GADD45A exon 2 (E) and MAZ exon 5 (F). Results are presented as percentage of input, and asterisks show significant changes (p-value < 0.05) compared to control (n = 6). **G-H)** ChRIP was performed in C33A cells expressing BRG1-wt with antibodies against Sam68, hnRNPU, and hnRNPL, and analysed with qPCR with the same primers as in Figure 2 for GADD45A exon 2 (G) and MAZ exon 5 (H). Results are presented as percentages, and asterisks show significant changes (p-value < 0.05) compared to control (n = 4).

Many RNA binding factors associate with both chromatin and the nascent RNA (Zhou et al., 2014; Chen et al., 2018) and we examined the association of factors with the mRNA of the genes by chromatin-RNA immunoprecipitation (ChRIP). HnRNPU, which was found at chromatin in MYL6 exon 6 in control cells, was also found to associate with the exon in the nascent RNA (Figure 5C). In BRG1 expressing cells, the binding of hnRNPU was lost on the nascent RNA, even though hnRNPU was associated with chromatin. Instead, SAM68, but not hnRNPL, bound to the exon in the nascent RNA (Figure 5C). SAM68 was not bound to a high level in chromatin, but possibly directly recruited to RNA by BRG1. No direct binding of BRG1 to exon 6 in RNA was detected, suggesting that it does not interact with RNA during transcription but rather associated with chromatin. In BRM expressing cells, hnRNPU was also excluded from the RNA, and no other interaction was detected (Supplementary S5C). In summary, BRG1 changed the interaction pattern of RNA binding proteins to the mRNA.

### BRG1 expression overcomes hnRNPU knock-down

To further investigate the role of the RNA binding factors on the splicing outcome of MYL6 exon 6 upon the expression of BRG1, we knocked down hnRNPL, hnRNPU and SAM68 using siRNAs (Supplementary Figure S5E). Knock-down of hnRNPL and hnRNPU in C33A control cells led to less inclusion of exon 6 in MYL6, while knock-down of SAM68 led to a significant increase in the inclusion rate (Figure 5D). Expression of BRG1 in cells in combination with hnRNPL knock-down did not restore the inclusion rate of exon 6, and the same level as in control cells was observed. SAM68 knock-down resulted in an even further increase of the inclusion rate in the presence of BRG1, suggesting that SAM68 has a fine-tuning role, dampening the increased rate in the presence of BRG1 (Figure 5D). Knock-down of hnRNPU in BRG1 expressing cells, however, restored the inclusion rate of MYL6 exon 6 to the level observed in BRG1 expressing cells in which hnRNPU was present. Knock-down of the factors in BRG1-mut expressing cells showed the same pattern of splicing; the splicing in hnRNPU knock-cells was restored by BRG1-mut expression to the level observed in BRG1-mut cells having hnRNPU (Supplementary Figure S5E). This suggests that hnRNPU promotes the inclusion of MYL6 exon 6 in control cells, but that BRG1 can replace this activity. BRG1 even enhanced the inclusion of the exon, possibly by allowing other RNA regulatory factors, such as SAM68, to interact with the RNA in the absence of hnRNPU.

### BRG1 alters the binding of RNA binding factors to GADD45A and MAZ

We also investigated how BRG1 and BRG-mut affected the recruitment of hnRNPU, hnRNPL and SAM68 to the affected exons in GADD45A and MAZ. BRG1 expression recruited the three factors to the alternative exon in GADD45A and MAZ cells, while the pattern of the factors in control cells differed. HnRNPL and SAM68 associated with GADD45A exon 2 in control cells, and in BRG1 and BRG1-mut expressing cells also hnRNPU was associated to the exon (Figure 5E and Supplementary Figure S5F). On the MAZ exon 5, only SAM68 associated in controls cells, and expression of BRG1 and BRG1-mut resulted in all three factors associating (Figure 5F and Supplementary Figure S5G).

The pattern of binding to the nascent RNA of the different factors on GADD45A exon 2 and MAZ exon 5 changed upon BRG1 expression, but in a different pattern to that found on MYL6 exon 6. SAM68 and hnRNPL, which were already present in chromatin without binding to RNA at GADD45A exon 2 in control cells, both bound to RNA in BRG1 expressing cells (Figure 5G). HnRNPU, which was associated with the exon upon BRG1 expression, did not bind to RNA (Figure 5G). The RNA binding pattern in MAZ provided a further variant; SAM68 associated with chromatin in control cells without binding to RNA, whereas hnRNPL associated only with the RNA. BRG1 expression changed the RNA binding pattern, only hnRNPU was associated with RNA, although all three factors associated with chromatin (Figure 5H). These results indicate that BRG1-SWI/SNF complexes modify the association of RNA binding factors with the exon and alter their interactions with the nascent RNA to fine-tune the splicing outcome.

## Discussion

mRNA alternative splicing is a combinatorial process, depending on a number of regulatory RNA binding proteins in addition to the general splicing machinery, chromatin states and transcription rate. Here, we demonstrate that the SWI/SNF complexes, which are mainly found at the promoter regulating transcription initiation (Masliah-Planchon et al., 2015; Clapier et al., 2017; Kadoch and Crabtree, 2015), influence alternative splicing by affecting the interaction of RNA binding proteins with chromatin and the nascent RNA at a subset of exons. Expression of the SWI/SNF ATPases BRG1 and BRM in the human SWI/SNF deficient cell line C33A promoted both exon inclusion and exon skipping, emphasising the complexity of splicing events: the splicing outcome depends on specific exon and intron features, as well as chromatin and RNA binding protein context. Previously, the ATPase BRM has been shown to be involved in alternative splicing of specific exons in mammalian cells, favouring inclusion in a process that does not require the ATPase activity (Batsché et al., 2006; Ito et al., 2008). The BRG1 protein binds RNA binding proteins, such as many proteins in snU2 and snU5 spliceosomes (Allemand et al., 2016; Yu et al., 2018), and participates in cleavage and polyadenylation (Yu et al., 2018). We show here that BRG1 is involved in splicing by both ATP-dependent and independent mechanisms, and that BRG1 and BRM can substitute for one another at some exons, but most targets are specific for each ATPase. The BRG1 ATPase independent exons displayed a high GC-content that were surrounded by high GC-content flanking regions compared to exons genome-wide, and no positioned nucleosome at the exon. This group resembled a group of exons defined in mammalian and avian genomes with a high GC-content, no differential GC-content in flanking regions, short introns, and no clear positioned nucleosome at the exon (Amit et al. 2012). This exon architecture is suggested to be identified by intron definition using splicing regulators (Amit et al., 2012; Gelfman et al., 2013).

Chromatin remodelling proteins play a role in alternative splicing by affecting the chromatin state. The nucleosome density (Tilgner et al., 2009; Luco et al., 2010; Zhou et al., 2014) and the histone modification state at exons (Alló et al., 2009; Tilgner et al., 2009; Luco et al., 2010; Enroth et al., 2012; Curado et al., 2015; Chen et al., 2018) have been proposed to define alternative exons for the splicing machinery and to affect the RNA-polymerase phosphorylation level and the transcription rate (Braunschweig et al., 2013; Fu and Ares, 2014; Zhou et al., 2014; Jonkers et al., 2014; Naftelberg et al., 2015; Fong et al., 2017; Nojima et al., 2018). The BRM affects alternative splicing by increasing the Ser5-P CTD state of RNA-polymerase II at alternative exons in HeLa cells (Batsché et al., 2006; Vorobyeva et al., 2012). In *Drosophila* BRM-SWI/SNF complexes have been proposed to use a different mechanism: it acts on stalled RNA pol II at a nucleosome block at alternative exons. SNR1 (an INI1/SNF5 orthologue) induces stalling, which prevents splicing, and the subsequent release is caused by remodelling of the nucleosome by BRM-SWI/SNF. This results in intron retention, without a change in the Ser5-P CTD during release and without a changed transcription rate. It was suggested that SWI/SNF operates by inhibiting splicing factors from binding to the RNA (Zraly and Dingwall, 2012). Our study of ATPase independent BRG1-SWI/SNF affected exons could not support any of the previously proposed mechanisms for SWI/SNF mediated splicing. Instead we suggest that the SWI/SNF complexes are involved in splicing using a different mechanism: fine-tuning the splicing outcome by influencing the binding of RNA binding proteins.

Many splicing regulatory factors and general splicing factors purify with SWI/SNF subunits (Zhao et al., 1998; Dellaire et al., 2002) and purifications of the snRNP U2 spliceosome component also include several SWI/SNF subunits (Makarov et al., 2012; Allemand et al., 2016). We performed an analysis of BRG1 and BRM interacting proteins in the RNP fraction (Yu et al., 2018) and it revealed that BRG1, in particular, interacted with U2 snRNP and U5-U6 snRNP factors that assemble early in the splicing cycle (Lardelli et al., 2010; Hegele et al., 2012; Agafonov et al., 2016; Haselbach et al., 2018). In addition to general splicing factors, we also found that BRG1 interacted with many regulatory RNA binding factors, such as hnRNPs and RNA helicases. Recent structural determinations of the spliceosome at different steps show that many rearrangements and compositional changes that occur during the splicing cycle require snRNA, splicing factors and regulatory factors (Bertram et al., 2017a; Bertram et al., 2017b;; Haselbach et al., 2018; Zhang et al., 2018). BRG1 recruits splicing factors and RNA binding proteins to affected exons, and we propose that it affects the interactions of RNA binding factors with their sites in RNA, which in turn influences the composition and the activity of the spliceosome and promote changes in splicing outcome.

The combinatorial nature of splicing regulation makes it is difficult to attribute changes in alternative splicing to only one RNA regulatory splicing factor. Instead, splicing can be achieved by several different mechanisms using the concurrent actions of a vast number of proteins and RNAs. Large-scale network analysis suggests that enhancer proteins interact by promoting the assembly of spliceosome sub complexes, whereas silencing occurs through RNA interactions (Ulrich and 2017; Guimarães et al., 2018). The combinatorial effect of different RNA splicing regulators is shown by the results of the knock-down of the factors on inclusion of MYL6 exon. MYL6 exon 6 depends on hnRNPL for inclusion, but without hnRNPL associating to the site (Hung et al., 2008; Rossbach et al., 2014; Vu et al., 2013; Shankarling et al., 2014; Cole et al., 2015). Our experiments show that hnRNPL is recruited to chromatin, but not RNA, in BRG1 expressing cells with higher inclusion observed. Instead, BRG1 excluded the hnRNPU, which was bound to RNA in control cells, from RNA, while BRG1 also promoted binding of the recruited SAM68to the RNA. In hnRNPU knock-down cells, BRG1 restored the higher inclusion level. This may be an effect of rearrangements in the spliceosome as a result of BRG1 rearrangement that allow different factors to bind to RNA. Combinatorial mechanisms have been proposed in which hnRNPs help to position splicing factors in the spliceosome and to block splice sites (Heinrich et al., 2009; Grillari et al., 2009; Howard et al., 2018). Other splicing factors, such as ZMAT2, fine-tune splicing by causing rearrangement to the spliceosome (Tanis et al., 2018). We speculate that chromatin remodelling factors also influence the spatial association of general splicing machinery with RNA; by recruiting, stabilising, and evicting splicing factors, to rearrange their interactions at the exons to fine-tune the splicing result (Figure 6).

**Figure 6.**
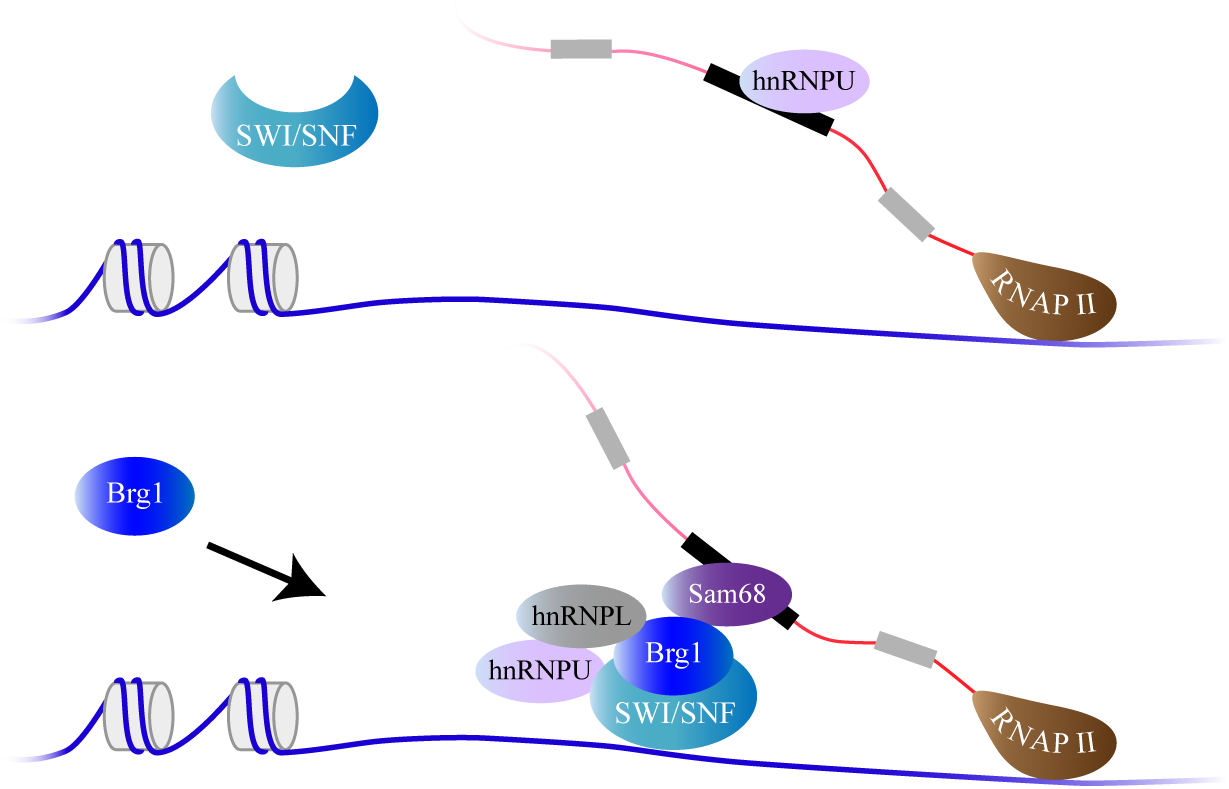
Model: BRG1 changes the recruitment pattern of RNA binding factor to alternative splicing exons in a context specific manner. MYL6 exon 6 has hnRNPU present in its vicinity at both chromatin and RNA level. Expression of BRG1 favours the recruitment of hnRNPL to this site in chromatin, while at the RNA level it recruits Sam68 and removes hnRNPU.

Dysregulated expressions of SWI/SNF components are often also found in malignant transformation, and may contribute to an altered gene expression that promotes cancer development (Biegel et al., 2014; Kadoch and Crabtree, 2015). Changes in splicing caused by mutations or deletions of snRNA and non-snRNP proteins are also tightly connected to malignant transformation. Our findings show that SWI/SNF complexes may contribute to cancer progression also by altering the splicing outcome. The functions of the different splice variants of the investigated genes have been associated with cancer transformation. MYL6 exon6 is more prevalent in smooth muscle, and the ratios of the splice variants are changed during cancer transformation, favouring a splice variant promoting migration (Li et al., 2006; Roberti et al., 2018). GADD45A also exhibits a changed splicing pattern in cancer cells, with the shorter splice form inhibiting cell cycle progression during stress (Zhang et al., 2009; Salvador et al., 2013; Carbonell et al., 2019). The longer MAZ exon 5 variant has been shown to inhibit the activation of inflammatory genes by the shorter MAZ isoform by binding more strongly to DNA (Ray et al., 2002; Triner et al., 2017). These examples indicate that SWI/SNF complexes affect the balances between splice variants with different functions, which may be an additional way that SWI/SNF targets are dysregulated during cancer progression.

A number of mechanisms operate to regulate the abundance of alternatively spliced exons. These mechanisms affect chromatin structure, transcription rate and the binding of splicing factor and RNA binding proteins. We show here that SWI/SNF complexes affect the splicing outcome by different mechanisms, both ATP-dependent and independent mechanisms, possibly determined by underlying exon features and recruited factors. SWI/SNF complexes have been shown to affect nucleosome structure and transcription rate and we propose an additional mechanism: BRG1 SWI/SNF complexes recruit RNA binding proteins and the general splicing machinery to alternative exons, and change the interaction of proteins in the nascent RNP to promote inclusion on exons with high GC content without a positioned nucleosome.

## Materials and methods

### Cell culture

Human HeLa and C33A cells (originally from ATCC) were cultured at 37°C and 5% CO2 in DMEM (HyClone) medium supplemented with 10% FBS, 50 U/ml penicillin and 50 μg/ml streptomycin.

### Exogenous expression and knock-down

C33A cells were transiently transfected for the expression of hBRG1 and the ATPase deficient BRG1 from the pBJ5-BRG1 and pBJ5-BRG1-K798R plasmids, respectively (Khavari et al., 1993). For hBRM and its ATPase deficient versions, pCG-hBrm and pCG-hBrm-K798R (Muchardt and Yaniv, 1993) were used. The pOPRSVI vector was used as control. Plasmids were transfected using Lipofectamin 2000 (Invitrogen) according to the manufacturer’s instructions for 48 hours before harvesting. BRG1 and BRM were knocked down in HeLa cells using siRNA. siRNA was transfected using RNAiMAX (Invitrogen) according to the manufacturer’s instructions. For knockdown experiments for hnRNPL, hnRNPU and SAM68, the same cells were transfected 24 hours after siRNA transfection with plasmids expressing BRG1, BRM and the mutated ATPases, and were incubated for an additional 48 hours before harvesting. SiRNAs for BRG1 (called SMARCA4) and BRM (called SMARCA2) and the RNA binding factors investigated are presented in Supplementary Table S6.

### RNA isolation and cDNA synthesis

RNA was extracted using Tri-reagent (Ambion/ThermoFisher) and treated with DNAse I (ThermoFisher). cDNA was synthesized with SuperScript III (Invitrogen/ThermoFisher) and oligodT according to the manufacturer’s instructions.

### qPCR

qPCR was performed in duplicate reactions using a KAPA SYBR Fast qPCR Kit (KAPABiosystem) in a QIAGEN Rotor-GeneQ system. Primers used are presented in Supplementary Table S5.

### RNA-seq, differential exon inclusion and gene expression analysis

Sequencing of 1 µg of RNA was performed with an Illumina HiSeq 2500, with 50 million reads depth. Reads were mapped with Tophat/2.0.4 to the Human genome assembly, build GRCh37. Gene counts were generated using HTseq/0.6.1 on bam files with duplicates included.

Alternative splicing was analyzed using MISO (Katz et al., 2010), and exons with a Bayes factor greater than 10 were considered to be differentially spliced. Exons showing opposite effects in the two replicates or in two different groups were discarded, as were exons with the same 5’ or 3’, and less than 50% of the length of the longest exon form. A given exon was only counted once, even if it was reported more than once in the MISO output. Differential gene expression was determined using DESeq2 with default parameters. C33A expressed exons were determined using FeatureCounts, and exons having a count in both replicates from pOPRSVI transfected cells were considered.

### Co-immunoprecipitation (Co-IP)

HeLa RNP extract was prepared as described in Tyagi et al. (2009). Briefly, nuclei were sonicated to obtain the chromatin fraction, and the chromatin was treated with RNAse A in PBS to release proteins bound to the nascent RNA (RNP fraction). The antibodies used were BRG1 antibody (Östlund Farrants et al., 1997), and the BRM and IgG antibody were from Abcam. Antibodies are presented in Supplementary Table S4.

### Immunoblotting blot

Cells were lysed in SDS-PAGE Laemli buffer containing 5% 2-mercaptoethanol. Protein extracts were separated by SDS-PAGE and transferred to a PVDF membrane (Millipore). Tubulin was used as a loading control (Abcam) for cell extracts and IgG antibody as negative control for co-IP. Antibodies against hnRNPL, DHX9, SAM68, SAP155 and THOC2 were from Abcam and listed in Table S4.

### Chromatin immunoprecipitation (ChIP)

C33A cells were fixed with 1% formaldehyde for 10 minutes at room temperature and chromatin extracted as described in Ryme et al. (2009). The chromatin was fragmented by sonication to fragments with a mean length of 500 bp. The antibodies used: SAM68, hnRNPL, hnRNPU, BAF155 and Ini1, were purchased from Abcam (Supplementary Table S4). Primers used in the analysis are presented in Supplementary Table S5.

### Chromatin RNA immunoprecipitation (ChRIP)

C33A cells were cross-linked by 1% formaldehyde-treated chromatin, and the RNA was extracted from the immunoprecipitated chromatin with Tri-reagent (Ambion/ThermoFisher), treated with DNAse I (ThermoFisher). cDNA was then synthesized with SuperScript III (Invitrogen/ThermoFisher) and random primers according to the manufacturer’s instructions. The antibodies used against SAM68, hnRNPL and hnRNPU were purchased from Abcam. Primers used in the analysis are presented in Supplementary Table S5.

### ChIP-seq data analysis

Published signal (bigwig files) from the ChIP-seq data was downloaded from ENCODE. Nucleosome positioning was downloaded from ENCODE/Stanford/BYU and histone modifications were downloaded from ENCODE/Broad. The files were transferred to the MISSISSIPPI Galaxy server (https://mississippi.snv.jussieu.fr/), lifted to hg19 reference genome when necessary using CrossMap (v0.2.7), and plotted using DeepTools2. The data for the plot were generated with computeMatrix 3.1.2, providing lists of included or skipped exons; 400 bp of exon and and 2000 bp upstream and downstream of those regions, 20 bp bin, missing values converted to 0 and mean selected as the statistic. The plot was generated with plotProfile (3.1.2) and used the “add standard error” mode.

### Chromatin states, regulatory regions and RNP motifs

The chromatin states of the SWI/SNF affected exons and all C33A exons expressed were determined with the intersection tool from the UCSC Table browser, using the Combined tables (ChromHMM+Segway) from the Genome Segments track. The intersections were counted and averaged for the six cell lines available. The regulatory regions were determined by intersecting the coordinates of affected exons with the oregannoAttr table from the UCSC ORegAnno track. RNP binding motifs present in SWI/SNF affected exons were determined using RBPmap (Paz et al., 2014) providing the exon coordinates and requesting all Human/Mouse motifs, with a high stringency level and conservation filter switched off.

### GC content

The sequences of affected exons and 500 bp upstream and downstream were retrieved with the Extract Genomic DNA tool (Galaxy Version 2.2.4). All sequences were aligned with transcription orientation and the average of C or G in each position was calculated. All exons were fitted in 100 bp: for this, the average of C or G was previously calculated for 1% of the total length of the exon. Smoothing was done with a moving average, factor 10 in Excel.

## Acknowledgements

The authors would like to acknowledge support from Science for Life Laboratory, the National Genomics Infrastructure (NGI) and Uppmax for providing assistance in massive parallel sequencing and computational infrastructure. SDM and MRF acknowledge funding from ERC Starting Grant (758397) and from the Strategic Research Area (SFO) programme of the Swedish Research Council through Stockholm University. NV acknowledges funding from The Swedish Research Council (2015–04553) and The Swedish Cancer Society (CAN 2016/460). AKÖF acknowledges funding from Stockholm University and Carl Tryggers Stiftelse för vetenskaplig forskning (CTS15:568). The funders had no role in study design, data collection and analysis, decision to publish, or preparation of the manuscript

---

**Supplementary Figure S1**

**A)** Number of exons with increased inclusion (blue) or skipped (orange) upon expression of exogenous BRG1, BRM, and their catalytically inactive versions. **B)** Number of exons that fall in all possible combinations between BRG1, BRG1-mut, BRM and BRM-mut. Coloured squares represent the presence in each single group. **C)** Venn diagram showing common genes affected by the expression of SWI/SNF ATPase subunits detected by DESeq2 (Love et al., 2014). **D)** The presence of SWI/SNF subunits in the genes of BRG1 affected exons was determined using publically available ChIP-seq data (Euskirchen et al., 2011). The presence of SWI/SNF subunits was considered positive if the peak interval overlapped at least 1 bp over the affected exon or the gene containing it. **E)** Transcription factor binding sites that overlapped BRG1 affected exons according to OREGANNO database. The figure shows 15 transcription factors with the highest numbers of sites associated to BRG1 affected exons. **F)** GC content (top panel) and nucleosome density (bottom panel) from ATPase dependent BRG1 affected exons that are also affected by BRM-wt exogenous expression +/- 500 bp harbouring regions divided into included (blue) and skipped (orange) exons. Exons and the harbouring +/-500 bp regions are shown. **G)** Prediction of the chromatin state at exons affected by BRG1 using publicly available data from chromatin in state segmentation by HMM from ENCODE. Top panel shows the state of all exons expressed in the C33A cells transfected with control plasmid pOPRSVI. Bottom panels show chromatin states for BRG1 affected exons according to whether they are BRG1 ATPase dependent (left) or ATPase independent (right), included (upper) or skipped (lower). Chromatin states were retrieved for the six cell lines available in the ENCODE database and the number of overlaps was averaged. An overlap was considered positive if the chromatin state interval overlapped at least 1 bp over the affected exon (shown as a percentage of the total amount of overlaps). **H)** Histone modification coverages around ATPase dependent BRG1 (left panels) and ATPase independent BRG1 affected exons (right panels). H3K4me3, H3K36me3, H3K9ac and H3K27ac data were obtained from publicly available data from the K562 cell line, at exons differentially included (blue) or skipped (orange) +/- 2 Kb harbouring region (light colour shows standard error).

**Supplementary Figure S2**

**A)** ChIP-qPCR shows association of BRG1 and BRM in the C33A cell line expressing inactive BRG1-mut to MYL6 exon 6 (left panel), GADD45A exon 2 (middle panel) and MAZ exon 5 (right panel). Significant changes (p-value < 0.05) compared to control are marked with asterisks (n = 6). **B)** ChIP-qPCR shows association of BRG1 and BRM in the C33A cell line expressing BRM or inactive BRM-mut to MYL6 exon 6 (left panel), GADD45A exon 2 (middle panel) and MAZ exon 5 (right panel). Significant changes (p-value < 0.05) compared to control are marked with asterisks (n = 6).

**Supplementary Figure S3**

**A-C)** ChIP-qPCR using antibodies against H3 (top panels) and H3K36me3 (bottom panels) targeting MYL6 gene (A), GADD45A exon 2 (B) and MAZ exon 5 (C) in C33A cells expressing BRM and BRM-mut. H3K36me3 levels were normalized to H3. The association is related to the association in control cells and significant values (p-value < 0.05) are denoted by asterisks (n = 3). **D-F)** ChIP-qPCR using antibodies against H3K4me3 (top panels), H3K9ac (middle panels) and H3K27ac (bottom panels) in C33A cells expressing BRG1, BRG1-mut, BRM and BRM-mut, targeting MYL6 exon 6 (D), GADD45A exon 2 (H) and MAZ exon 5 (I). The signal from histone modifications was first normalized to the H3 signal for each treatment, and the association is related to the association in control cells. Asterisks denote significant differences (p-value < 0.05) compared to control (n = 3). **G-I)** ChIP-qPCR using antibodies against polymerase II CTD (top panels), and phosphorylated serines 2 and 5 from polymerase II CTD (P-CTD, bottom panels) targeting MYL6 gene (A), GADD45A exon 2 (B) and MAZ exon 5 (C) in C33A cells expressing BRM1 and BRM-mut. The association is related to the association in control cells and significant values (p-value < 0.05) are denoted by asterisks (n = 5).

**Supplementary Figure S4**

**A)** Most enriched GO terms from BRG1 and BRM interacting proteins related to splicing, represented in –log10(p-value). The GO Ontology database, released on 9 August 2018was used, and the PANTHER Overrepresentation Test (released on 5 December 2017). **B)** Western blot from co-IP performed with BRG1 and BRM antibodies in the RNP fraction of HeLa cells. Membranes were probed with antibodies against proteins detected in the MASS spectroscopy data from Yu et al., 2018. IgG is used as negative control. **C)** Western blot from co-IP using antibodies against Sam68 and membranes probed with BRG1 and BRM antibodies. IgG is used as negative control.

**Supplementary Figure S5**

**A)** ChIP-qPCR targeting MYL6 exon 6 in C33A cells expressing BRM and BRM-mut was performed in C33A cells expressing BRM-wt and BRM-mut with antibodies against Sam68, hnRNPU, hnRNPL, DHX15, SYF1, THOC2, hnRNPA1 and hnRNP2B1. Significant changes (p-value < 0.05) compared to control are marked with asterisks. **B)** ChIP was performed in C33A cells expressing BRM-wt with antibodies against Sam68, hnRNPU, hnRNPL and analysed with qPCR with the same primers as in Figure 2 for MYL6 promoter, exon 6 and exon 7. Results are presented as percentage of input and asterisks show significant changes (p-value < 0.05) compared to control (n = 6). **C)** ChRIP was performed in C33A cells expressing BRM-wt with antibodies against Sam68, hnRNPU, and hnRNPL, and analysed with qPCR with the same primers as in Figure 2 for MYL6 exon 6. Results are presented as percentages and asterisks show significant changes (p-value < 0.05) compared to control (n = 4). **D)** Immunoblots showing the efficiency of knock-downs of RNA binding factor in C33A cells, 3 days after transfection of siRNA. Tubulin was used as loading control. The size in KDa is marked at the right. **E)** Changes in the inclusion of SWI/SNF affected exons after knocking down Sam68, hnRNPU and hnNRPL. C33A cells were transfected with siRNA targeting splice regulatory factors interacting with BRG1 and BRM, incubated for 24h, and transfected with BRG1-mut for an additional 48 h. The primers presented in Figure 2A were used to assess exon 6 in MYL6. Values were normalised to scr siRNA for each ATPase expressed, and asterisks show significant changes (p-value < 0.05) compared to control (n = 3). **F and G)** ChIP was performed in C33A cells expressing BRG1-mut with antibodies against Sam68, hnRNPU, and hnRNPL, and analysed with qPCR with the same primers as in Figure 2 for GADD45A exon 2 and MAZ exon 5. Results are presented as percentage of input and asterisks show significant changes (p-value < 0.05) compared to control (n = 6).

**Supplementary Table S1**

Exons affected by expression of the SWI/SNF ATPases in C33A cells. The exon that were differentially expressed were identified by the MISO algorithm.

**Supplementary Table S2**

Differentially expressed genes in C33A cells expressing the ATPases.

**Supplementary Table S3**

Genes in which both the expression and exons were affected by the expression of SWI/SNF ATPases.

**Supplementary Table S4**

Sequences of the SiRNAs used in the study.

**Supplement Table S5**

Antibodies used in the study.

**Supplementary Table S6**

Primer pairs used for qPCR in the study.

## References

Agafonov DE, van Santen M, Kastner B, Dube P, Will CL, Urlaub H, Lührmann R. ATPγS stalls splicing after B complex formation but prior to spliceosome activation. RNA. 2016 Sep;22(9):1329–37. doi: 10.1261/rna.057810.116.

Allemand E, Myers MP, Garcia-Bernardo J, Harel-Bellan A, Krainer AR, Muchardt C. A Broad Set of Chromatin Factors Influences Splicing. PLoS Genet. 2016 Sep23;12(9):e1006318. doi: 10.1371/journal.pgen.1006318.

Alló M, Buggiano V, Fededa JP, Petrillo E, Schor I, de la Mata M, Agirre E, Plass M, Eyras E, Elela SA, Klinck R, Chabot B, Kornblihtt AR. Control of alternative splicing through siRNA-mediated transcriptional gene silencing. Nat Struct Mol Biol. 2009 Jul;16(7):717–24. doi: 10.1038/nsmb.1620.

Alló M, Agirre E, Bessonov S, Bertucci P, Gómez Acuña L, Buggiano V, Bellora N, Singh B, Petrillo E, Blaustein M, Miñana B, Dujardin G, Pozzi B, Pelisch F, Bechara E, Agafonov DE, Srebrow A, Lührmann R, Valcárcel J, Eyras E, Kornblihtt AR. Argonaute-1 binds transcriptional enhancers and controls constitutive and alternative splicing in human cells. Proc Natl Acad Sci U S A. 2014 Nov 4;111(44):15622–9. doi: 10.1073/pnas.1416858111.

Ameur A, Zaghlool A, Halvardson J, Wetterbom A, Gyllensten U, Cavelier L, Feuk L. Total RNA sequencing reveals nascent transcription and widespread co-transcriptional splicing in the human brain. Nat Struct Mol Biol. 2011 Nov 6;18(12):1435–40. doi: 10.1038/nsmb.2143.

Ameyar-Zazoua M, Rachez C, Souidi M, Robin P, Fritsch L, Young R, Morozova N, Fenouil R, Descostes N, Andrau JC, Mathieu J, Hamiche A, Ait-Si-Ali S, Muchardt C, Batsché E, Harel-Bellan A. Argonaute proteins couple chromatin silencing to alternative splicing. Nat Struct Mol Biol. 2012 Oct;19(10):998–1004. doi: 10.1038/nsmb.2373.

Amit M, Donyo M, Hollander D, Goren A, Kim E, Gelfman S, Lev-Maor G, Burstein D, Schwartz S, Postolsky B, Pupko T, Ast G. Differential GC content between exons and introns establishes distinct strategies of splice-site recognition. Cell Rep.2012 May 31;1(5):543–56. doi: 10.1016/j.celrep.2012.03.013.

Batsché E, Yaniv M, Muchardt C. The human SWI/SNF subunit Brm is a regulator of alternative splicing. Nat Struct Mol Biol. 2006 Jan;13(1):22–9.

Bertram K, Agafonov DE, Liu WT, Dybkov O, Will CL, Hartmuth K, Urlaub H, Kastner B, Stark H, Lührmann R. Cryo-EM structure of a human spliceosome activated for step 2 of splicing. Nature. 2017 Feb 16;542(7641):318–323. doi: 10.1038/nature21079.

Bertram K, Agafonov DE, Dybkov O, Haselbach D, Leelaram MN, Will CL, Urlaub H, Kastner B, Lührmann R, Stark H. Cryo-EM Structure of a Pre-catalytic Human Spliceosome Primed for Activation. Cell. 2017 Aug 10;170(4):701–713.e11. doi: 10.1016/j.cell.2017.07.011.

Biegel JA, Busse TM, Weissman BE. SWI/SNF chromatin remodeling complexes and cancer. Am J Med Genet C Semin Med Genet. 2014 Sep;166C(3):350–66. doi:10.1002/ajmg.c.31410.

Braunschweig U, Gueroussov S, Plocik AM, Graveley BR, Blencowe BJ. Dynamic integration of splicing within gene regulatory pathways. Cell. 2013 Mar 14;152(6):1252–69. doi: 10.1016/j.cell.2013.02.034.

Carbonell C, Ulsamer A, Vivori C, Papasaikas P, Böttcher R, Joaquin M, Miñana B, Tejedor JR, de Nadal E, Valcárcel J, Posas F. Functional Network Analysis Reveals the Relevance of SKIIP in the Regulation of Alternative Splicing by p38 SAPK. Cell Rep. 2019 Apr 16;27(3):847–859.e6. doi: 10.1016/j.celrep.2019.03.060.

Chen W, Feng P, Ding H, Lin H. Classifying Included and Excluded Exons in Exon Skipping Event Using Histone Modifications. Front Genet. 2018 Oct 1;9:433. doi: 10.3389/fgene.2018.00433.

Cho EJ, Takagi T, Moore CR, Buratowski S. mRNA a capping enzyme is recruited to the transcription complex by phosphorylation of the RNApolymerase II carboxy-terminal domain. Genes Dev. 1997 Dec 15; 11(24):3319–26.

Clapier CR, Iwasa J, Cairns BR, Peterson CL. Mechanisms of action and regulation of ATP-dependent chromatin-remodelling complexes. Nat Rev Mol Cell Biol. 2017 Jul;18(7):407–422. doi: 10.1038/nrm.2017.26.

Cole BS, Tapescu I, Allon SJ, Mallory MJ, Qiu J, Lake RJ, Fan HY, Fu XD, Lynch KW. Global analysis of physical and functional RNA targets of hnRNP L reveals distinct sequence and epigenetic features of repressed and enhanced exons. RNA. 2015 Dec;21(12):2053–66. doi: 10.1261/rna.052969.115.

Curado J, Iannone C, Tilgner H, Valcárcel J, Guigó R. Promoter-like epigenetic signatures in exons displaying cell type-specific splicing. Genome Biol. 2015 Oct23;16:236. doi: 10.1186/s13059-015-0797-8. Erratum in: Genome Biol.2016;17(1):52.

Custódio N, Carmo-Fonseca M. Co-transcriptional splicing and the CTD code. Crit Rev Biochem Mol Biol. 2016 Sep;51(5):395–411.

De Conti L, Baralle M, Buratti E. Exon and intron definition in pre-mRNA splicing.Wiley Interdiscip Rev RNA. 2013 Jan-Feb;4(1):49–60. doi: 10.1002/wrna.1140.

Decristofaro MF, Betz BL, Rorie CJ, Reisman DN, Wang W, Weissman BE. Characterization of SWI/SNF protein expression in human breast cancer cell lines and other malignancies. J Cell Physiol. 2001 Jan;186(1):136–45.

Dellaire G, Makarov EM, Cowger JJ, Longman D, Sutherland HG, Lührmann R, Torchia J, Bickmore WA. Mammalian PRP4 kinase copurifies and interacts with components of both the U5 snRNP and the N-CoR deacetylase complexes. Mol Cell Biol. 2002 Jul;22(14):5141–56.

Dujardin G, Lafaille C, de la Mata M, Marasco LE, Muñoz MJ, Le Jossic-Corcos C, Corcos L, Kornblihtt AR. How slow RNA polymerase II elongation favors alternative exon skipping. Mol Cell. 2014 May 22;54(4):683–90. doi: 10.1016/j.molcel.2014.03.044

Di Giammartino DC, Nishida K, Manley JL. Mechanisms and consequences of alternative polyadenylation. Mol Cell. 2011 Sep 16;43(6):853–66. doi:10.1016/j.molcel.2011.08.017.

Dyvinge H. Regulation of alternative mRNA splicing: old players and new perspectives. FEBS Lett. 2018 Sep;592(17):2987–3006. doi: 10.1002/1873-3468.13119.

Enroth S, Bornelöv S, Wadelius C, Komorowski J. Combinations of histone modifications mark exon inclusion levels. PLoS One. 2012;7(1):e29911. doi: 10.1371/journal.pone.0029911.

Euskirchen GM, Auerbach RK, Davidov E, Gianoulis TA, Zhong G, Rozowsky J, Bhardwaj N, Gerstein MB, Snyder M. Diverse roles and interactions of the SWI/SNF chromatin remodeling complex revealed using global approaches. PLoS Genet. 2011 Mar;7(3):e1002008. doi: 10.1371/journal.pgen.1002008.

Fong N, Saldi T, Sheridan RM, Cortazar MA, Bentley DL. RNA Pol II Dynamics Modulate Co-transcriptional Chromatin Modification, CTD Phosphorylation, and Transcriptional Direction. Mol Cell. 2017 May 18;66(4):546–557.e3. doi: 10.1016/j.molcel.2017.04.016.

Fontana GA, Rigamonti A, Lenzken SC, Filosa G, Alvarez R, Calogero R, Bianchi ME, Barabino SM. Oxidative stress controls the choice of alternative last exons via a Brahma-BRCA1-CstF pathway. Nucleic Acids Res. 2017 Jan 25;45(2):902–914. doi: 10.1093/nar/gkw780.

Fu XD, Ares M Jr. Context-dependent control of alternative splicing by RNA-binding proteins. Nat Rev Genet. 2014 Oct;15(10):689–701. doi: 10.1038/nrg3778.

Garavís M, González-Polo N, Allepuz-Fuster P, Louro JA, Fernández-Tornero C, Calvo O. Sub1 contacts the RNA polymerase II stalk to modulate mRNA synthesis. Nucleic Acids Res. 2017 Mar 17;45(5):2458–2471. doi: 10.1093/nar/gkw1206.

Gelfman S, Cohen N, Yearim A, Ast G. DNA-methylation effect on cotranscriptional splicing is dependent on GC architecture of the exon-intron structure. Genome Res. 2013 May;23(5):789–99. doi: 10.1101/gr.143503.112.

Georgomanolis T, Sofiadis K, Papantonis A. Cutting a Long Intron Short: Recursive Splicing and Its Implications. Front Physiol. 2016 Nov 29;7:598.

Grillari J, Löscher M, Denegri M, Lee K, Fortschegger K, Eisenhaber F, Ajuh P, Lamond AI, Katinger H, Grillari-Voglauer R. Blom7alpha is a novel heterogeneous nuclear ribonucleoprotein K homology domain protein involved in pre-mRNA splicing that interacts with SNEVPrp19-Pso4. J Biol Chem. 2009 Oct 16;284(42):29193–204. doi: 10.1074/jbc.M109.036632.

Guimarães PR Jr, Pires MM, Cantor M, Coltri PP. Interaction paths promote module integration and network-level robustness of spliceosome to cascading effects. Sci Rep. 2018 Nov 28;8(1):17441. doi: 10.1038/s41598-018-35160-6.

Gunderson FQ, Johnson, TL. Acetylation by the transcriptional coactivator Gcn5 plays a novel role in co-transcriptional spliceosome assembly. PLoS Genet., 2009 5, e1000682.

Hargreaves DC, Crabtree GR. ATP-dependent chromatin remodeling: genetics, genomics and mechanisms. Cell Res. 2011 Mar; 21(3): 396–420. Published online 2011 Mar 1. doi: 10.1038/cr.2011.32

Harlen KM, Trotta KL, Smith EE, Mosaheb MM, Fuchs SM, Churchman LS. Comprehensive RNA Polymerase II Interactomes Reveal Distinct and Varied Roles for Each Phospho-CTD Residue. Cell Rep. 2016 Jun 7;15(10):2147–2158. doi: 10.1016/j.celrep.2016.05.010

Haselbach D, Komarov I, Agafonov DE, Hartmuth K, Graf B, Dybkov O, Urlaub H, Kastner B, Lührmann R, Stark H. Structure and Conformational Dynamics of the Human Spliceosomal B^act^ Complex. Cell. 2018 Jan 25;172(3):454–464.e11. doi: 10.1016/j.cell.2018.01.010.

Hegele A, Kamburov A, Grossmann A, Sourlis C, Wowro S, Weimann M, Will CL, Pena V, Lührmann R, Stelzl U. Dynamic protein-protein interaction wiring of the human spliceosome. Mol Cell. 2012 Feb 24;45(4):567–80. doi:10.1016/j.molcel.2011.12.034.

Heinrich B, Zhang Z, Raitskin O, Hiller M, Benderska N, Hartmann AM, Bracco L, Elliott D, Ben-Ari S, Soreq H, Sperling J, Sperling R, Stamm S. Heterogeneous nuclear ribonucleoprotein G regulates splice site selection by binding to CC(A/C)-rich regions in pre-mRNA. J Biol Chem. 2009 May 22;284(21):14303–15. doi: 10.1074/jbc.M901026200.

Hirose Y, Ohkuma Y. Phosphorylation of the C-terminal domain of RNA polymerase II plays central roles in the integrated events of eucaryotic gene expression. J Biochem. 2007 May; 141(5):601–8.

Hnilicova J, Hozeifi S, Duskova E, Icha J., Tomankova T Stanek D. Histone deacetylase activity modulates alternative splicing. PLoS One, 2011 6, e16727.

Hollander D, Naftelberg S, Lev-Maor G, Kornblihtt AR, Ast G. How Are Short Exons Flanked by Long Introns Defined and Committed to Splicing? Trends Genet. 2016 Oct;32(10):596–606. doi: 10.1016/j.tig.2016.07.003.

Hota SK, Bruneau BG. ATP-dependent chromatin remodeling during mammalian development. Development. 2016 Aug 15;143(16):2882–97. doi: 10.1242/dev.128892.

Howard JM, Lin H, Wallace AJ, Kim G, Draper JM, Haeussler M, Katzman S, Toloue M, Liu Y, Sanford JR. HNRNPA1 promotes recognition of splice site decoys by U2AF2 in vivo. Genome Res. 2018 May;28(5):689–698. doi: 10.1101/gr.229062.117.

Hou Y, Huang H, Hu W, Sun X. Histone modification influence skipped exons inclusion. Journal of Bioinformatics and Computational Biology 2017 Vol. 15, No. 1750003 DOI: 10.1142/S0219720017500032

Hsin JP Manley JL. The RNA polymerase II CTD coordinates transcription and RNA processing. Genes Dev. 2012 26, 2119–2137.

Hung LH, Heiner M, Hui J, Schreiner S, Benes V, Bindereif A. Diverse roles of hnRNP L in mammalian mRNA processing: a combined microarray and RNAi analysis. RNA. 2008 Feb;14(2):284–96.

Iannone C, Valcárcel J. Chromatin’s thread to alternative splicing regulation. Chromosoma. 2013 Dec;122(6):465–74. doi: 10.1007/s00412-013-0425-x.

Ip JY, Schmidt D, Pan Q, Ramani AK, Fraser AG, Odom DT, Blencowe BJ. Global impact of RNA polymerase II elongation inhibition on alternative splicing regulation. Genome Res. 2011 Mar;21(3):390–401. doi: 10.1101/gr.111070.110.

Ito T, Watanabe H, Yamamichi N, Kondo S, Tando T, Haraguchi T, Mizutani T, Sakurai K, Fujita S, Izumi T, Isobe T, Iba H. Brm transactivates the telomerase reverse transcriptase (TERT) gene and modulates the splicing patterns of its transcripts in concert with p54(nrb). Biochem J. 2008 Apr 1;411(1):201–9.

Jordán-Pla A, Yu S, Waldholm J, Källman T, Östlund Farrants AK, Visa N. SWI/SNF regulates half of its targets without the need of ATP-driven nucleosome remodeling by Brahma. BMC Genomics. 2018 May 18;19(1):367. doi: 10.1186/s12864-018-4746-2.

Jonkers I, Kwak H, Lis JT. Genome-wide dynamics of Pol II elongation and its interplay with promoter proximal pausing, chromatin, and exons. Elife. 2014 Apr 29;3:e02407. doi: 10.7554/eLife.02407.

Katz Y, Wang ET, Airoldi EM, Burge CB. Analysis and design of RNA sequencing experiments for identifying isoform regulation. Nat Methods. 2010 Dec;7(12):1009–15. doi: 10.1038/nmeth.1528.

Kadoch C, Crabtree GR. Mammalian SWI/SNF chromatin remodeling complexes and cancer: Mechanistic insights gained from human genomics. Sci Adv. 2015 Jun12;1(5):e1500447. doi: 10.1126/sciadv.1500447.

Kim YE, Park C, Kim KE, Kim KK. Histone and RNA-binding protein interaction creates crosstalk network for regulation of alternative splicing. Biochem Biophys Res Commun. 2018 Apr 30;499(1):30–36. doi: 10.1016/j.bbrc.2018.03.101.

Kornblihtt AR. Coupling transcription and alternative splicing. Adv Exp MedBiol. 2007;623:175–89.

Kornblihtt AR, Schor IE, Allo M, Blencowe BJ. When chromatin meets splicing. Nat Struct Mol Biol. 2009 Sep;16(9):902–3.

Lardelli RM, Thompson JX, Yates JR 3rd, Stevens SW. Release of SF3 from the intron branchpoint activates the first step of pre-mRNA splicing. RNA. 2010 Mar;16(3):516–28. doi: 10.1261/rna.2030510.

Lee Y, Rio DC. Mechanisms and Regulation of Alternative Pre-mRNA Splicing. Annu Rev Biochem. 2015;84:291–323. doi: 10.1146/annurev-biochem-060614-034316.

Lesurf R, Cotto KC, Wang G, Griffith M, Kasaian K, Jones SJ, Montgomery SB, Griffith OL; Open Regulatory Annotation Consortium. ORegAnno 3.0: a community-driven resource for curated regulatory annotation. Nucleic Acids Res. 2016 Jan 4;44(D1):D126–32. doi: 10.1093/nar/gkv1203.

Li X, Kazan H, Lipshitz HD, Morris QD. Finding the target sites of RNA-binding proteins. Wiley Interdiscip Rev RNA. 2014 Jan-Feb; 5(1):111–30.

Luco RF, Pan Q, Tominaga K, Blencowe BJ, Pereira-Smith OM, Misteli T. Regulation of alternative splicing by histone modifications. Science. 2010 Feb19;327(5968):996–1000. doi: 10.1126/science.1184208.

Love MI, Huber W, Anders S. Moderated estimation of fold change and dispersion for RNA-seq data with DESeq2. Genome Biol. 2014;15(12):550.

Makarov EM, Owen N, Bottrill A, Makarova OV. Functional mammalian spliceosomal complex E contains SMN complex proteins in addition to U1 and U2 snRNPs. Nucleic Acids Res. 2012; 40: 2639–2652. doi: 10.1093/nar/gkr1056

Masliah-Planchon J, Bièche I, Guinebretière JM, Bourdeaut F, Delattre O.SWI/SNF chromatin remodeling and human malignancies. Annu Rev Pathol.2015;10:145–71. doi: 10.1146/annurev-pathol-012414-040445.

McCracken S, Fong N, Rosonina E, Yankulov K, Brothers G, Siderovski D, Hessel A, Foster S, Shuman S, Bentley DL. 5’-Capping enzymes are targeted to pre-mRNA by binding to the phosphorylated carboxy-terminal domain of RNA polymerase II. Genes Dev. 1997 Dec 15;11(24):3306–18.

Moteki S, Price D. Functional coupling of capping and transcription of mRNA. Mol Cell. 2002 Sep;10(3):599–609.

Muchardt C, Yaniv M. A human homologue of Saccharomyces cerevisiae SNF2/SWI2and Drosophila brm genes potentiates transcriptional activation by the glucocorticoid receptor. EMBO J. 1993 Nov;12(11):4279–90.

Naftelberg S, Schor IE, Ast G, Kornblihtt AR. Regulation of alternative splicing through coupling with transcription and chromatin structure. Annu RevBiochem. 2015;84:165–98. doi: 10.1146/annurev-biochem-060614-034242

Nojima T, Rebelo K, Gomes T, Grosso AR, Proudfoot NJ, Carmo-Fonseca M. RNA Polymerase II Phosphorylated on CTD Serine 5 Interacts with the Spliceosome during Co-transcriptional Splicing. Mol Cell. 2018 Oct 18;72(2):369–379.e4. doi: 10.1016/j.molcel.2018.09.004.

Obrdlik A, Kukalev A, Louvet E, Farrants AK, Caputo L, Percipalle P. The histone acetyltransferase PCAF associates with actin and hnRNP U for RNA polymerase II transcription. Mol Cell Biol. 2008 Oct;28(20):6342–57. doi: 10.1128/MCB.00766-08.

Östlund Farrants AK, Blomquist P, Kwon H, Wrange O. Glucocorticoid receptor-glucocorticoid response element binding stimulates nucleosome disruption by the SWI/SNF complex. Mol Cell Biol. 1997 Feb;17(2):895–905.

Pradeepa, MM, Sutherland HG, Ule J., Grimes G.R Bickmore WA. Psip1/Ledgf p52 Binds Methylated Histone H3K36 and Splicing Factors and Contributes to the Regulation of Alternative Splicing. PLoS Genet., 2012 8, e1002717.

Ray BK, Murphy R, Ray P, Ray A. SAF-2, a splice variant of SAF-1, acts as a negative regulator of transcription. J Biol Chem. 2002 Nov 29;277(48):46822–30.

Rossbach O, Hung LH, Khrameeva E, Schreiner S, König J, Curk T, Zupan B, Ule J, Gelfand MS, Bindereif A. Crosslinking-immunoprecipitation (iCLIP) analysis reveals global regulatory roles of hnRNP L. RNA Biol. 2014;11(2):146–55. doi: 10.4161/rna.27991.

Ryme J, Asp P, Böhm S, Cavellán E, Farrants AK. Variations in the composition of mammalian SWI/SNF chromatin remodelling complexes. J Cell Biochem. 2009 Oct 15;108(3):565–76. doi: 10.1002/jcb.22288.

Saldi T, Cortazar MA, Sheridan RM, Bentley DL. Coupling of RNA Polymerase II Transcription Elongation with Pre-mRNA Splicing. J Mol Biol. 2016 Jun 19;428(12):2623–2635. doi: 10.1016/j.jmb.2016.04.017.

Saldi T, Fong N, Bentley DL. Transcription elongation rate affects nascent histone pre-mRNA folding and 3’ end processing. Genes Dev. 2018 Feb 1;32(3-4):297–308. doi: 10.1101/gad.310896.117.

Salvador JM, Brown-Clay JD, Fornace AJ Jr. Gadd45 in stress signaling, cell cycle control, and apoptosis. Adv Exp Med Biol. 2013;793:1–19. doi: 10.1007/978-1-4614-8289-5_1.

Schwartz S, Ast G. Chromatin density and splicing destiny: on the cross-talk between chromatin structure and splicing. EMBO J. 2010 May 19;29(10):1629–36. doi: 10.1038/emboj.2010.71.

Shankarling G, Cole BS, Mallory MJ, Lynch KW. Transcriptome-wide RNA interaction profiling reveals physical and functional targets of hnRNP Lin human T cells. Mol Cell Biol. 2014 Jan;34(1):71–83. doi: 10.1128/MCB.00740-13.

Shukla S, Oberdoerffer S. Co-transcriptional regulation of alternative pre-mRNA splicing. Biochim Biophys Acta. 2012 Jul;1819(7):673–83. doi: 10.1016/j.bbagrm.2012.01.014.

Shenasa H, Hertel KJ. Combinatorial regulation of alternative splicing. Biochim Biophys Acta Gene Regul Mech. 2019 Jul 2. pii: S1874-9399(19)30102-6. doi: 10.1016/j.bbagrm.2019.06.003.

Sims RJ 3rd, Millhouse S, Chen CF, Lewis BA, Erdjument-Bromage H, Tempst P, Manley JL, Reinberg D. Recognition of trimethylated histone H3 lysine 4 facilitates the recruitment of transcription postinitiation factors and pre-mRNA splicing. Mol Cell. 2007 Nov 30;28(4):665–76.

Spain MM. and Govind, C.K. A role for phosphorylated Pol II CTD in modulating transcription coupled histone dynamics. Transcription 2011 2, 78–81.

Tanis SEJ, Jansen PWTC, Zhou H, van Heeringen SJ, Vermeulen M, Kretz M, Mulder KW. Splicing and Chromatin Factors Jointly Regulate Epidermal Differentiation. Cell Rep. 2018 Oct 30;25(5):1292–1303.e5. doi: 10.1016/j.celrep.2018.10.017.

Tilgner H, Nikolaou C, Althammer S, Sammeth M, Beato M, Valcárcel J, Guigó R. Nucleosome positioning as a determinant of exon recognition. Nat Struct Mol Biol.2009 Sep;16(9):996–1001. doi: 10.1038/nsmb.1658.

Tilgner H, Knowles DG, Johnson R, Davis CA, Chakrabortty S, Djebali S, Curado J, Snyder M, Gingeras TR, Guigó R. Deep sequencing of subcellular RNA fractions shows splicing to be predominantly co-transcriptional in the human genome but inefficient for lncRNAs. Genome Res. 2012 Sep;22(9):1616–25. doi: 10.1101/gr.134445.111.

Triner D, Castillo C, Hakim JB, Xue X, Greenson JK, Nuñez G, Chen GY, Colacino JA, Shah YM. Myc-Associated Zinc Finger Protein Regulates the Proinflammatory Response in Colitis and Colon Cancer via STAT3 Signaling.Mol Cell Biol. 2018 Oct 29;38(22). pii: e00386–18. doi: 10.1128/MCB.00386-18.

Tyagi A, Ryme J, Brodin D, Ostlund Farrants AK, Visa N. SWI/SNF associates with nascent pre-mRNPs and regulates alternative pre-mRNA processing. PLoS Genet. 2009 May;5(5):e1000470. doi: 10.1371/journal.pgen.1000470.

Ulrich, A.K.C., and Wahl, M.C. (2017). Human MFAP1 is a cryptic ortholog of the Saccharomyces cerevisiae Spp381 splicing factor. BMC Evol. Biol. 17, 91

Vu NT, Park MA, Shultz JC, Goehe RW, Hoeferlin LA, Shultz MD, Smith SA, Lynch KW, Chalfant CE. hnRNP U enhances caspase-9 splicing and is modulated by AKT-dependent phosphorylation of hnRNP L. J Biol Chem. 2013 Mar 22;288(12):8575–84. doi: 10.1074/jbc.M112.443333.

Vizlin-Hodzic D, Runnberg R, Ryme J, Simonsson S, Simonsson T. SAF-A forms a complex with BRG1 and both components are required for RNA polymerase II mediated transcription. PLoS One. 2011;6(12):e28049. doi: 10.1371/journal.pone.0028049.

Waldholm J, Wang Z, Brodin D, Tyagi A, Yu S, Theopold U, Farrants AK, Visa N. SWI/SNF regulates the alternative processing of a specific subset of pre-mRNAs in *Drosophila* melanogaster. BMC Mol Biol. 2011 Nov 2;12:46. doi:10.1186/1471-2199-12-46.

Wang ET, Sandberg R, Luo S, Khrebtukova I, Zhang L, Mayr C, Kingsmore SF, Schroth GP, Burge CB. Alternative isoform regulation in human tissue transcriptomes. Nature. 2008 Nov 27;456(7221):470–6. doi: 10.1038/nature07509.

Witten JT, Ule J. Understanding splicing regulation through RNA splicing maps.Trends Genet. 2011 Mar;27(3):89–97. doi: 10.1016/j.tig.2010.12.001.

Wong AK, Shanahan F, Chen Y, Lian L, Ha P, Hendricks K, Ghaffari S, Iliev D, Penn B, Woodland AM, Smith R, Salada G, Carillo A, Laity K, Gupte J, Swedlund B, Tavtigian SV, Teng DH, Lees E. BRG1, a component of the SWI-SNF complex, is mutated in multiple human tumor cell lines. Cancer Res. 2000 Nov 1;60(21):6171–7.

Vorobyeva NE, Nikolenko JV, Nabirochkina EN, Krasnov AN, Shidlovskii YV, Georgieva SG. SAYP and Brahma are important for ‘repressive’ and ‘transient’ Pol II pausing. Nucleic Acids Res. 2012 Aug;40(15):7319–31. doi: 10.1093/nar/gks472.

Yearim A, Gelfman S, Shayevitch R, Melcer S, Glaich O, Mallm JP, Nissim-Rafinia M, Cohen AH, Rippe K, Meshorer E, Ast G. HP1 is involved in regulating the global impact of DNA methylation on alternative splicing. Cell Rep. 2015 Feb 24;10(7):1122–34. doi: 10.1016/j.celrep.2015.01.038.

Yu S, Jordán-Pla A, Gañez-Zapater A, Jain S, Rolicka A, Östlund Farrants AK, Visa N. SWI/SNF interacts with cleavage and polyadenylation factors and facilitates pre-mRNA 3’ end processing. Nucleic Acids Res. 2018 May 31. doi:10.1093/nar/gky438.

Zhang Y, Beezhold K, Castranova V, Shi X, Chen F. Characterization of an alternatively spliced GADD45alpha, GADD45alpha1 isoform, in arsenic-treated epithelial cells. Mol Carcinog. 2009 May;48(5):454–64. doi: 10.1002/mc.20483.

Zhang X, Yan C, Zhan X, Li L, Lei J, Shi Y. Structure of the human activated spliceosome in three conformational states. Cell Res. 2018 Mar;28(3):307–322. doi: 10.1038/cr.2018.14.

Zhang J, Kuo CC, Chen L. GC content around splice sites affects splicing through pre-mRNA secondary structures. BMC Genomics. 2011 Jan 31;12:90. doi: 10.1186/1471-2164-12-90.

Zhao K, Wang W, Rando OJ, Xue Y, Swiderek K, Kuo A, Crabtree GR. Rapid and phosphoinositol-dependent binding of the SWI/SNF-like BAF complex to chromatin after T lymphocyte receptor signaling. Cell. 1998 Nov 25;95(5):625–36.

Zhou HL, Luo G, Wise JA, Lou H. Regulation of alternative splicing by local histone modifications: potential roles for RNA-guided mechanisms. Nucleic Acids Res. 2014 Jan; 42(2):701–13.

Zraly CB, Dingwall AK. The chromatin remodeling and mRNA splicing functions ofthe Brahma (SWI/SNF) complex are mediated by the SNR1/SNF5 regulatory subunit.Nucleic Acids Res. 2012 Jul;40(13):5975–87. doi: 10.1093/nar/gks288.

